# Large Scale Cell Painting Guided Compound Selection Reveals Activity Cliffs and Functional Relationships

**DOI:** 10.1101/2025.05.16.654292

**Authors:** Maxime Sanchez, Nicolas Bourriez, Ihab Bendidi, Ethan Cohen, Ivan Svatko, Elaine Del Nery, Hamza Tajmouati, Guillaume Bollot, Laurence Calzone, Auguste Genovesio

## Abstract

Traditional structure-based pre-screen compound selection relies on the assumption that chemical similarity implies similar biological activity. This paradigm narrows the exploration of chemical space and often fails to account for functional convergence, where structurally diverse compounds act through distinct targets to produce similar phenotypic effects. As a result, compounds with therapeutic potential may be overlooked. To overcome this constraint, we introduce a training-free, transfer learning-based method for large scale compound preselection that leverages deep phenotypic profiling of human cells. Notably, this enables robust pairwise comparison of phenotypic signatures across any source of the entire JUMP-CP, the largest publicly available cell painting dataset (112,480 compounds), preserving biological signals while mitigating batch effects. Validated across 65 high-throughput assays—including in vitro and in cellulo systems—our method provides efficient pre-screen enrichment of biologically active compounds, bypassing the blind spots of structure-centric approaches. Interestingly, because it is large scale, it also allows for a comprehensive analysis of structure–phenotypic activity relationships, revealing potentially thousands of compound activity cliffs, where minimal chemical changes in structure may result in profound phenotypic shifts. We show that these cliffs capture subtle, atom-level determinants of bioactivity that cannot be accessed by structure-based models. Furthermore, we demonstrate that structurally diverse compounds targeting different genes in the same biological pathway can induce either convergent or opposite phenotypes—a phenomenon validated across 30 pathways, hundreds of genes, and thousands of compounds. Finally, to support the broader community, we propose Phenoseeker, a web-based tool enabling instant retrieval of JUMP-CP compounds with similar phenotypic profiles. Together, these findings position phenotypic profiling not merely as a complementary tool, but as a transformative and scalable framework for navigating chemical space through a biological lens. By capturing rich morphological signatures that reflect functional outcomes—regardless of structural similarity—this approach enables the discovery of bioactive compounds, novel mechanisms of action, and unexpected target-pathway relationships. Applied at the scale of the JUMP-CP dataset, phenotypic profiling emerges as a powerful strategy for prioritizing compounds, illuminating activity cliffs, and accelerating the identification of therapeutically relevant candidates across diverse biological contexts.

## Introduction

Over the past decades, phenotypic drug discovery (PDD) has re-emerged as a powerful strategy for identifying novel therapeutic agents ^1,2^. In contrast to target-based approaches that focus solely on predetermined molecular targets, such as specific proteins, PDD harnesses observable cellular or organismal phenotypes ^2^. Although both strategies have produced effective therapeutics, growing evidence indicates that PDD is more successful at delivering first-in-class drugs ^3^. Its target-agnostic nature is especially advantageous for addressing polygenic diseases or conditions linked to undruggable targets. Moreover, PDD offers the simultaneous assessment of many potential targets and the filtering of compounds based on key criteria such as solubility, toxicity, and cellular accessibility ^4^. It also entails drawbacks including the need for subsequent target deconvolution and the challenge of deciphering off-target interactions from direct effects. Importantly, the success of a drug discovery campaign highly depends on the relevance of the compound library, which, in the case of PDD, is hard to predict. In fact, an efficient pre-screening compound selection method for PDD should select compounds that are likely to reproduce the desired phenotype in the considered assay.

A relevant approach toward this goal is the Cell Painting assay ^5^, a standardized, multiplexed imaging technique introduced in 2013 ^6^. This method produces high-content, high-resolution images that capture compound -or genetically-induced cellular changes by labeling non specific organelles (including nuclei, nucleoli, mitochondria, actin, and tubulin networks) chosen for their generic nature, unrelated to any specific phenotype. As a result, it provides an unbiased, holistic view of the cell state. Alone or in combination with other omics data, Cell Painting has helped to decipher cell death pathways ^7^, to assess chemical exposures in non-tumorigenic breast cells ^8^, to study gene roles in glaucoma ^9^, or to predict drug-induced toxicity in muscle cells^10^. Initially applied to U2OS cells, the assay has now been adapted for various cell lines^11^, live imaging^12^, human induced pluripotent stem cells (hiPSC)^13^, and even 3D cell spheroids^14^, demonstrating its impact in both fundamental research and therapeutic discovery^15^.

In parallel, the resurgence of deep learning^16^ has revolutionized biotechnology, as highlighted by AlphaFold’s breakthrough in protein structure prediction^17^. Researchers have integrated ideas and architectures from diverse deep learning areas, such as representation learning^18^, multimodal learning^19^, and generative models^20^ to reveal more nuanced and distinct phenotypic patterns in cell imaging data. This progress has led to novel approaches, such as InfoAlign^21^, Cell Painting CNN^22^, or CLOOME^23^, which can predict molecular properties or drug mechanisms of action ^25,29,33,34^. It has even helped uncover biases in CRISPR-Cas9 genome editing^24^ and has been applied in genetic perturbation studies ^25,26^. In short, deep learning models can be used to convert microscopy images of cells to a vector called a *deep phenotypic profile* which can further be exploited to compute similarities between phenotypic responses to various perturbations.

If the hypothesis is made that two compound perturbations producing the same cell phenotype are likely to impact the same function, then phenotypic profiles computed from Cell Painting images could be used as a proxy to perform an efficient compound library pre-selection. We recently validated this hypothesis with a small proof of concept on 3 screens using a first Cell Painting screen of 10,000 compounds ^27,28^. Other works have used cell phenotypes to identify candidates most likely to exhibit a desired bioactivity (e.g., protein inhibition, cell death, or pathway activation)^27,29–37^. Recent methods using supervised classifiers require hundreds of compounds with known bioactivities^37^. Some works use handcrafted cell features (sometimes these features are combined with chemical structures ^36^ or genetic profiles), while other approaches use unimodal ^30,31,33^ or multimodal^29^ deep encoders. Altogether, these methods require a training procedure, (i.e., training data), and thus are not easily transferable to drug discovery projects. A transfer learning strategy such as the one we proposed in Cohen et al^27^ does not necessitate any training step and is straightforward to use as it only requires a positive control reference. However, this preselection has been so far limited in scale due to the unavailability of large cell painting datasets.

Scaling our pre-selection method to a much larger library is now within reach, thanks to the introduction of the JUMP-CP dataset in 2023^38^, which contains images of over 112,000 unique compound perturbations. However, for this massive dataset to be assembled, it was co-produced by 12 laboratories around the world, spread on various experimental batches and acquired with different microscopes. Consequently, it exhibits considerable non-biological variability, known as “*batch effects*”^39^. Altogether scaling compound selection to a large library dispatched on multiple sites and multiple batches such as JUMP-CP is not straightforward and it is yet unclear what combination of deep encoder, normalization strategy and profile replicate aggregation would be optimal at this scale to produce phenotypic profiles robust enough for an efficient compound selection. For this reason, to our knowledge, there is no work to date that benefits from an exhaustive view on the dataset compound perturbations for this task.

In this work, we identified an effective encoding-normalization-aggregation strategy to make possible robust comparison of any couple of compounds through their phenotypic profiles across the entire JUMP-CP dataset. We then devised a training-free transfer learning approach to perform an efficient preselection of compounds prior to a screen. We thoroughly validated its efficiency across 65 diverse screens, from internal and public databases of high-throughput assays, for a total of 40k+ readouts, covering both *in vitro* and *in cellulo* bioactivities. Interestingly, we identified that our method performs large “jumps” in the vast chemical space (10³³ drug-like compounds^40^) using phenotypic similarity as a biological proxy. These results show a greater chemical diversity among the selected compounds confirming preliminary observations made using a small Cell Painting dataset from one of the twelve JUMP-CP laboratories^37^. We further demonstrated its ability to uncover activity cliffs that are typically missed by traditional structure-based selection approaches^41^. Our findings also show that compounds targeting the same signaling pathway tend to induce related phenotypic profiles, a trend we validated on 30 pathways spanning hundreds of genes and thousands of compounds. In order to facilitate real-world applications and accelerate drug discovery, we created a website (https://www.phenoseeker.bio.ens.psl.eu/) where researchers can retrieve in one click a large preselection from the 112,480 JUMP-CP compounds that are most likely to exhibit a bioactivity that is similar to the positive control query.

## Results

### Building phenotypic profiles that enhances signal and reduces batch effect over the whole JUMP-CP

Being able to compare cell phenotypes induced by all the compound perturbations across the whole JUMP-CP dataset required several processing steps (encoding, aggregating and normalizing). However, no clear state-of-the-art method has yet emerged for the whole JUMP-CP. We have developed a flexible pipeline (**Figure 1a**) that includes various approaches for each of these steps, some of which can be found in previous works ^25,39^. Our goal was to identify the best combination that would minimize batch effects while preserving biologically relevant signals. The first step, extracting a feature vector from multiple 5-channel images acquired using various microscope-imaging configurations or plate formats (i.e., 384 or 1536 wells), can be achieved either using hand-crafted methods (e.g., CellProfiler^42^) or various deep-learning encoders. These encoders may be pretrained on natural images (e.g., ResNet50^43^, DINOv2^44^) or on microscopy images (e.g., ChAda-ViT^45^, OpenPhenom^46^) (see <u>Methods</u>). Following this, the feature vectors of all fields of view from one well need to be aggregated in order to obtain one profile per experimental sample. Wells distributed over different plates in different batches from different laboratories then need to be aligned through a normalization step. For instance, one can use mock treatment control units in each batch to center and scale features or perform a Typical Variation Normalization (TVN)^47^, which aligns both first-order statistics and covariance structures. Other methods from the field of transcriptomics, such as neural network–based *scVI*^48^, mixture-model *Harmony*^49^, or nearest neighbor *Scanorama*^50^, as well as dedicated training losses ^51,52^ and deep-learning architectures^51^ were considered, but these necessitate a training step which we aimed to avoid for robustness. Finally, replicate wells need to be aggregated into a single feature profile per condition referred to as “phenotypic profile”.

**Figure 1:**
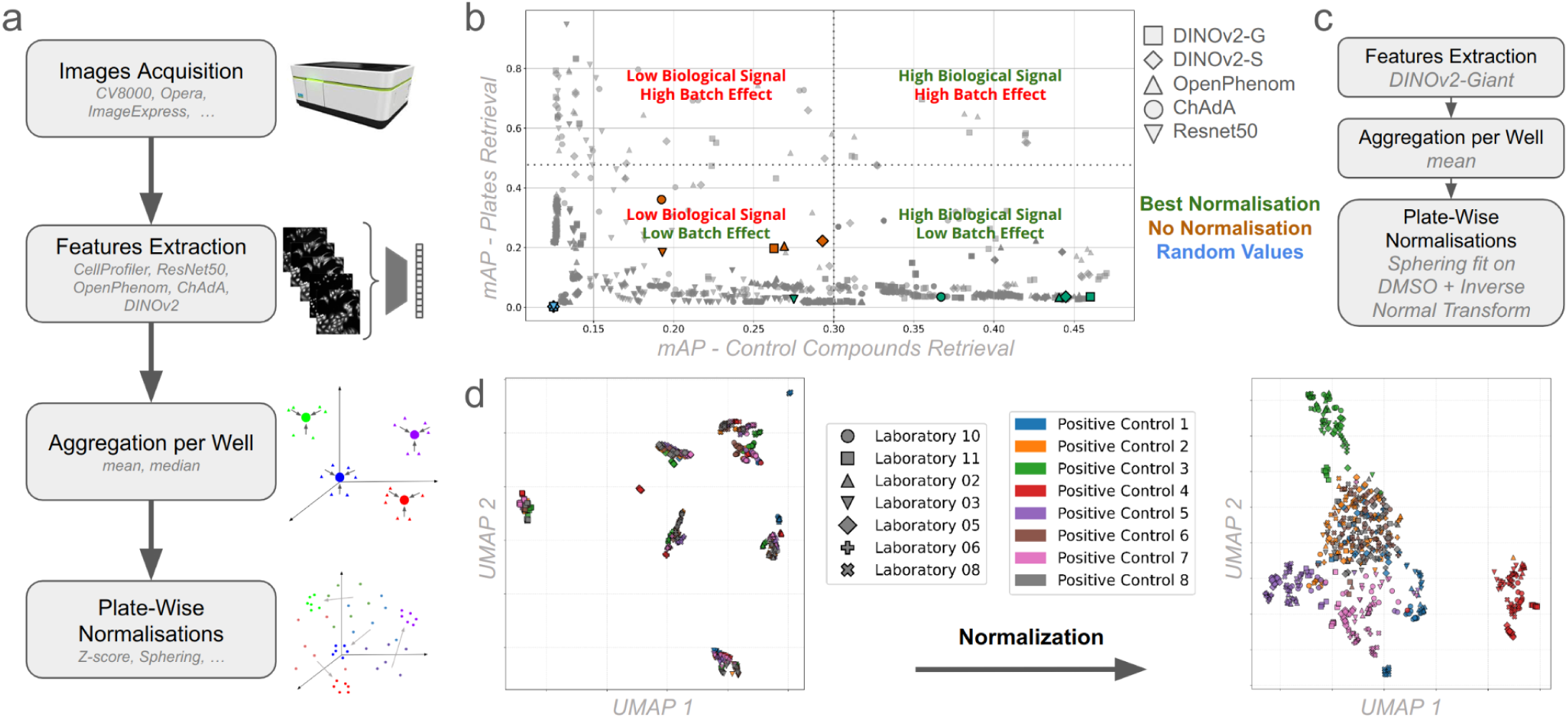
Generating robust Phenotypic Profiles from JUMP-CP Cell Painting images. **(a)** Typical pipeline steps from image acquisition to normalized well profiles. At each of these steps (feature extraction, aggregation and normalization), various methods can be chosen **(b)** We performed a sweep of possible pipelines using control well replicate samples from JUMP-CP. For each possible pipeline (represented on this plot by a single point), after processing all samples, we checked how well the pipeline retrieved sample replicates from the same perturbation (X axis, the higher the better) and could confound replicate samples of negative control whatever the experimental plate (Y axis, the lower the better) by computing two mean Average Precisions (mAP) values. The former evaluates the preservation of biological signals while the latter assesses effectiveness in correcting batch effects. **(c)** Scheme of the selected processing of Cell Painting images **(d)** UMAP^54^ Visualization of Well Profiles Across Laboratories. UMAP^54^ plots display well phenotypic profiles from 27 plates (only control wells) across 9 laboratories, before (left) and after (right) normalization, showing improved integration of phenotypic profiles.

To identify which combination of these steps best preserves biological signal while mitigating batch effects, two metrics were used (**Figure 1b**). The first metric captures the preservation of biological signal, as measured by the mean average precision (mAP) for eight positive control compounds present on every experimental plate. We seek to obtain a high value since it indicates that profiles induced by the same compound treatment are close in phenotypic space. The second metric captures batch effect removal as measured by the mAP on experimental plate labels (see <u>Methods</u>). We seek to approach the low value of a random retrieval indicating that profiles from the same plate (therefore from the same batch and laboratory) are not particularly closer than those from different plates. By performing a thorough sweep of more than 5000 combinations (see <u>Methods</u>) of processing steps described above, we identified a relevant combination of steps that optimize our criteria. First, we determined that DINOv2-Giant provides a better encoding than other deep encoders (including those trained on microscopy images) or handcrafted solutions such as CellProfiler^42^ (**Figure 1b**). Following this, we determined that the most effective normalization approach consisted in fitting a whitening matrix to the DMSO negative control wells of each plate, then applying it to all the sample wells of this plate ^53^ then finally applying an inverse normal transformation (INT) to ensure each feature follows a normal distribution across the plate (**Figure 1**). Before normalization, wells from the same laboratory tend to cluster together, reflecting strong batch effects and making phenotypic profile comparison across experimental batches irrelevant. After normalization, these clusters become more intermixed while, on the contrary, positive control replicates across laboratories cluster together, indicating a reduction in batch variation and an enhancement of biological signal over experimental noise (**Figure 1d**). This step enables the comparison of any pair of samples, regardless of their origin, thus making the JUMP-CP large compound library fully usable for further applications.

### Large-scale Cell Painting-based selection boosts active compound yield

By leveraging on this representation that allows comparisons across a large range of perturbations, we propose a practical compound selection method using similarity of Cell Painting phenotypic profiles as a proxy. Indeed, we hypothesized that two different compound perturbations exhibiting the same phenotypic response shared similar bioactivity. Our pipeline then uses cosine similarity (see <u>Methods</u>) to compare the phenotypic profiles of 112,480 compounds in JUMP-CP to the positive control (**Figure 2a**), a compound that is known to induce the desired bioactivity. In the following, we name the cosine similarity between two phenotypic profiles of compounds the “phenotypic similarity” between those two compounds. This approach enables rapid and efficient selection of bioactive compounds without requiring any machine learning training or database of compounds with known bioactivities. Designed for simplicity and usability, our method allows any researcher to identify compounds with a desired bioactivity using only a single positive control. If the phenotypic profile of this positive control is not already available in the JUMP-CP dataset, it can be generated by imaging the control with the Cell Painting assay and computing a normalized phenotypic profile with our pipeline. We validated our method using 65 screens, some of which were previously performed at the BioPhoenix platform at Institut Curie (16) and others were collected from ChEMBL (49) open source data, as well as 5 targets from Lit-PCBA benchmark ^55^ (see <u>Methods</u>). To evaluate the quality of our compound selection for these screens, we employed the normalized enrichment factor (nEF), defined as the enrichment factor (EF) divided by the maximum possible EF (**Figure 2a**, <u>Methods</u>). As shown in **Figure 2b**, c, d, nEF values were calculated for the selection of the top 5% compounds with the highest phenotypic similarity to the positive control. We observed comparable results at other selection percentages (see **Supplementary Fig. 4**), and the raw EF values are provided in **Supplementary Fig. 14**. Because the compounds tested in the JUMP-CP dataset and in each evaluation screen only partially overlapped, we lacked perfect positive controls for each of them. To address this, we systematically treated each identified hit from Institut Curie, ChEMBL and Lit-PCBA screens (i.e., bioactive compound) as a potential positive control, calculating the nEF for each of them. We then considered two evaluation scenarios: a pessimistic scenario where the nEF is averaged over all hits, thus accounting for false positives or weaker hits (**Figure 2b**, c, d, green bars); and an optimistic scenario where the highest nEF obtained was considered, reflecting a more realistic configuration where the positive control is a tool compound with a strong phenotype and a well-characterized bioactivity (**Figure 2b**, c, d, blue bars). In most tested screens, selecting the top 5% compounds most similar to the positive control, under the pessimistic scenario, outperformed (or sometimes matched) the hit rate of the original selection (**Figure 2b**, c, d, purple bars). Importantly, our method almost never yields a worse hit ratio than the original screens, even when suboptimal or false positive compounds were taken into account. Moreover, when a highly effective positive control is considered (the optimistic scenario), our method always drastically improves hit enrichment, occasionally achieving the best possible selection of compounds and, at the very least, consistently doubling the proportion of active compounds compared to the initial selection. Notably, the method’s strong performance on 5 target proteins from the Lit-PCBA benchmark—which is specifically designed to assess target-dependent effects—demonstrates that it does not merely capture false positives or off-target signals, a common pitfall in phenotypic drug discovery. This target-specific validation confirms that our approach reliably enriches the library for compounds with relevant bioactivity, thereby enhancing its practical utility in drug discovery.

**Figure 2:**
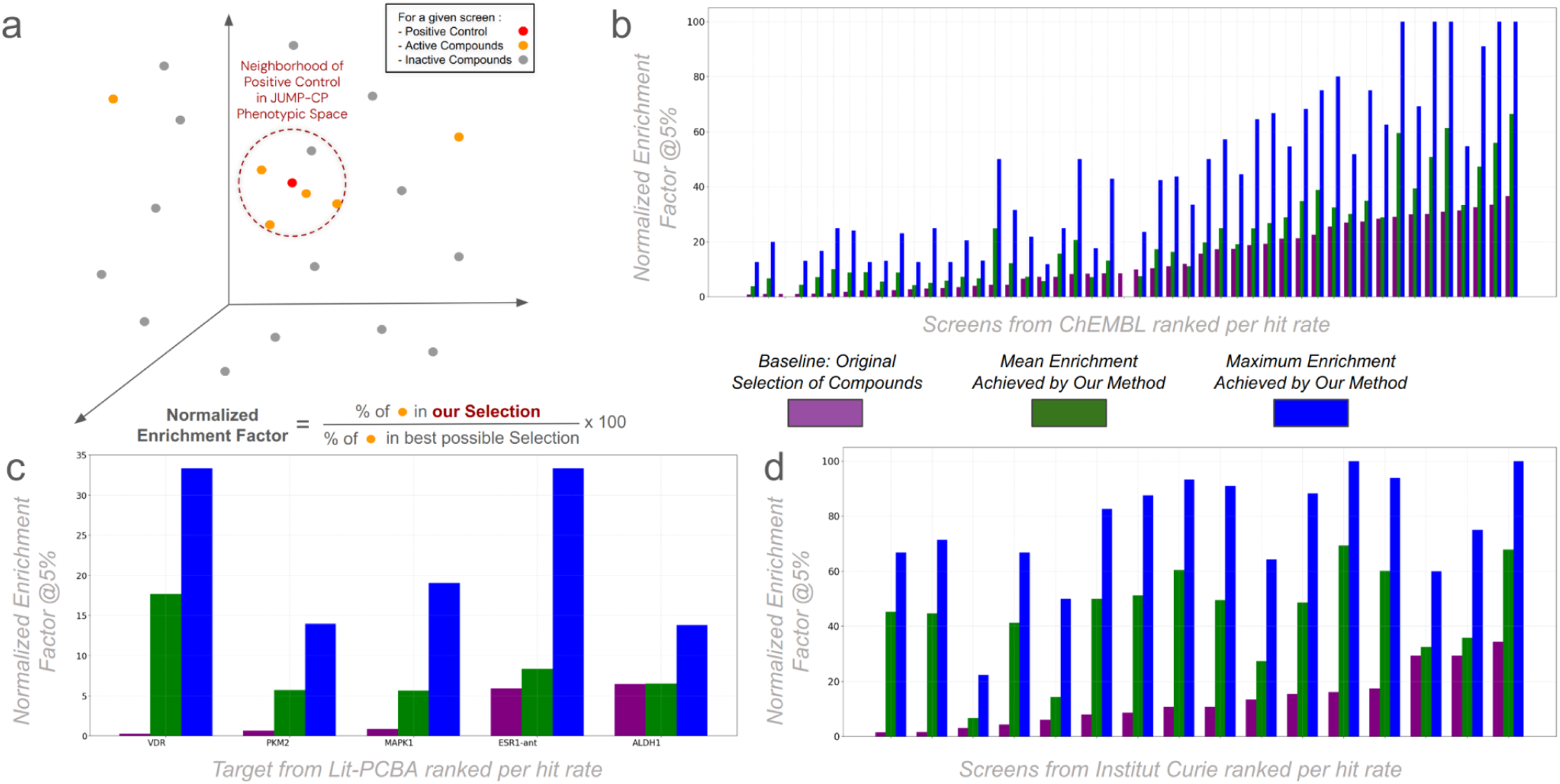
Library selection through robust JUMP-CP profiles boost screening hit rates. **(a)** Schematic of our selection method where the top k% closest JUMP-CP neighbors (dotted sphere) of a screen positive control (red dot) are selected for screening. To evaluate the capability of this selection process to increase screening hit rate, we applied it to past screens and thus computed the fraction of hits that fall into this selection (orange dots). **(b)(c)(d)** Evaluation of our method across diverse screens and sources. Mean (green bars) and maximum (blue bars) normalized Enrichment Factor (nEF) are displayed, comparing our method with the original selection (purple bar). There are 49 screens from ChEMBL **(b)**, 5 targets from Lit-PCBA **(c)**, and 16 screens from the Curie Screening Platform **(d)**. nEF is determined based on the top 5% compounds with the highest phenotypic similarity to the positive control.

### Phenotypic-based compound selection increases the chemical diversity of biologically active compounds

We further investigated the structural diversity of the compounds selected by our approach. To encode the structural topology of a chemical compound, we use Morgan fingerprints, a type of circular molecular descriptor widely used in cheminformatics^56^. Using the Tanimoto similarity^57^, we measured the structural similarity between the positive control from each of the Institut Curie and ChEMBL screens and the compounds selected through our biological proxy (see <u>Methods</u>). This structural similarity remained consistently low (**Figure 3a**) and was statistically significantly lower than the structural similarity between the positive control and the top 5% compounds most similar in structure (see <u>Methods</u> and **Supplementary Table 2** for statistical details). Remarkably, the structural similarity between the positive control and the entire pool of screened compounds was comparable to that between the positive control and the top 5% of compounds with the highest phenotypic similarity (**Figure 3a**). We extended this finding for each compound of the entire JUMP-CP dataset by computing the Tanimoto similarity between its top 5% with highest phenotypic similarity and its top 5% most structurally similar (**Figure 3a**). To illustrate this, we generated UMAP^54^ embeddings of both phenotypic profiles and Morgan fingerprints (**Figure 3b**) for compounds tested in a given cell-based fluorescence assay at the BioPhoenix platform. The positive control (in red) and the compounds selected with our method (in blue) appear in distinct regions of chemical space, further illustrating our approach’s capacity to identify chemically diverse compounds that may share similar bioactivities. Our method identifies compounds with different chemical scaffolds that produce phenotypic profiles similar to the positive control (**Figure 3c**). This broadens the chemical diversity of selected compounds, and therefore future hits, and creates opportunities to discover compounds with potentially improved ADME-Toxicity profiles or fewer intellectual property constraints. Conversely, we observed instances where compounds with highly similar structures produced very different phenotypes (**Figure 3d**).

**Figure 3:**
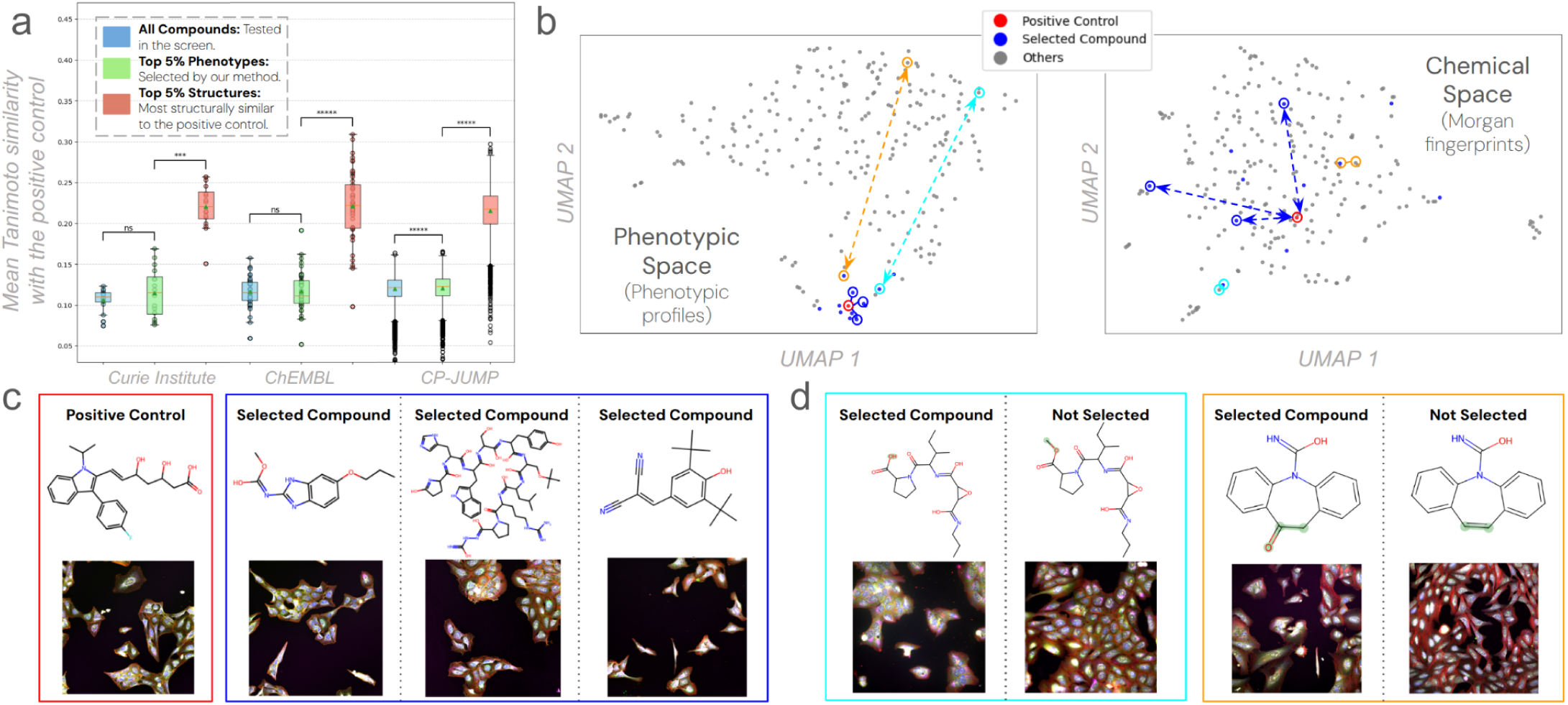
Selection through similar Cell Painting profiles yield a variety of compound structures. **(a)** For illustration, the Mean Tanimoto structure similarity between the positive control of a screen from Institut Curie and three groups of compounds: all compounds tested in the screen (blue), the top 5% phenotypically similar compounds (green), and the top 5% structurally similar compounds (red). The same computation is then performed for a screen from ChEMBL and the whole JUMP-CP using each compound as a positive control. **(b)** On the left, *UMAP*^54^ *of the phenotypic profiles* for compounds tested in the screen from Institut Curie. On the right, *UMAP of the structural fingerprints* for the same compounds. Positive control is shown in red; the 5% phenotypically closest compounds are shown in blue. Arrows point to illustration compounds from **(c)** and **(d)**. **(c)** *Examples of selected compounds* (close to the positive control in phenotypic space but distant in structural space). **(d)** *Examples of pairs of compounds* that are close in structural space but distant in phenotypic space. For each compound, chemical structures and Cell Painting images are shown.

### Clustering analogs per phenotype reveals activity cliffs

The last results motivated us to explore whether the JUMP-CP dataset could also reveal structural relationships between compounds explaining variation in their phenotypic profiles. Specifically, we aimed to examine whether compounds that share the same core scaffold, even with minor chemical variations, can produce distinct phenotypic profiles, denoting “phenotypic activity cliffs.” An activity cliff is a phenomenon in which two structurally similar compounds show a significant difference in biological activity.

First, compounds from the JUMP-CP dataset were grouped according to their Demis-Murcko scaffold^58^, resulting in 532 distinct chemical series with 6 compounds or more (**Figure 4a**, <u>Methods</u>). Then, in each of these series, compounds were split in two clusters of phenotypes (defined as the highest level of a hierarchical clustering of their phenotypic profiles) and the intra- and inter-cluster phenotypic and structural similarities were computed (**Figure 4a**). 81 series (2277 compounds in total, see **Supplementary Fig. 12** for examples of scaffolds) showed a substantially higher intra-cluster phenotypic similarity compared to their inter-cluster phenotypic similarity (**Figure 4c**, <u>Methods</u>) while showing minimal differences structure-wise by the same metrics (**Figure 4d**) thus defining potentially thousands of compounds activity cliffs. We repeated this experiment using hierarchical clustering with 3, 4, and 5 clusters, consistently obtaining similar results (see **Supplementary Fig. 11**). This finding suggests that phenotypic profiles reveal activity cliffs that are difficult to predict by quantitative structure-activity relationship (QSAR) methods^41^. This illustrates the advantage of PDD and the value of using Cell Painting phenotypes to complement the structural description of compounds, enhancing chemical space exploration in drug discovery projects.

**Figure 4:**
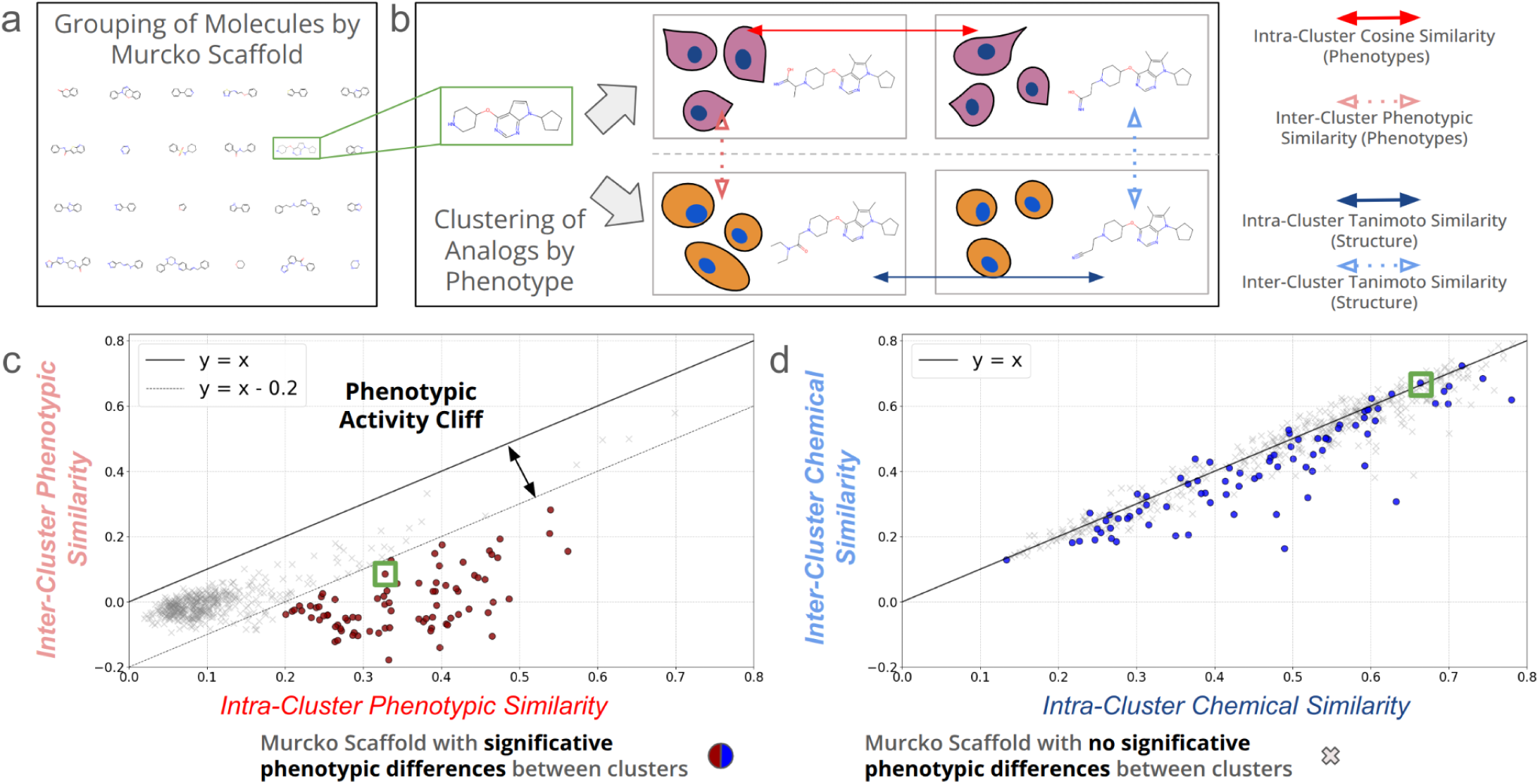
Systematic identification of activity cliffs. **(a)**. JUMP-CP compounds were grouped by Murcko scaffold. **(b)** For each group of compounds, a hierarchical clustering based on their phenotypic profiles was performed. Phenotypic (cosine, red) and Structural (Tanimoto, blue) similarity within and between these clusters were computed. **(c)** Intra-versus inter-cluster phenotypic similarity. 81 scaffold-based groups exposing a substantially higher intra-than inter-cluster phenotypic similarity are highlighted in red. **(d)** Intra-versus inter-cluster structural similarities. The 81 scaffolds highlighted in red on panel c are highlighted in blue here. The green squares represent the example of the chemical series illustrated in (b).

### Clustering analogs per phenotype reveals key chemical functions

To expand on our previous finding, we cherry-picked examples of chemical series where compounds exhibit distinct phenotypic profiles (**Figure 5**, **Supplementary Fig. 9**, **Supplementary Fig. 8**, **Supplementary Fig. 10**). For instance, a cluster of such a series was made of three compounds (C1, C2, C3 **Figure 5b**) killing the cells (**Figure 5a**). Interestingly, they share three specific moieties: (1) a bromine atom, (2) a methyl group, and (3) a sp3 carbon chain linked to the nitrogen atom (**Figure 5b**, in green). These three compounds are also significantly different from the negative DMSO control (**Figure 5a**). In contrast, compounds C4 through C11 induce different phenotypes, both from C1–C3 and from one another (**Figure 5c**). These differences may stem from the protonation of the scaffold’s nitrogen (green highlight), as well as other structural variations (purple highlights). For instance, compound C9, which is the Murcko scaffold reference of this series, induces a phenotype similar to DMSO (**Figure 5a**), indicating no significant phenotypic effect. Meanwhile, adding different chemical substitutions to the Murcko scaffold induces subtle changes in the cellular phenotype, yet these differences remain clearly distinguishable with our phenotypic profiles.

**Figure 5:**
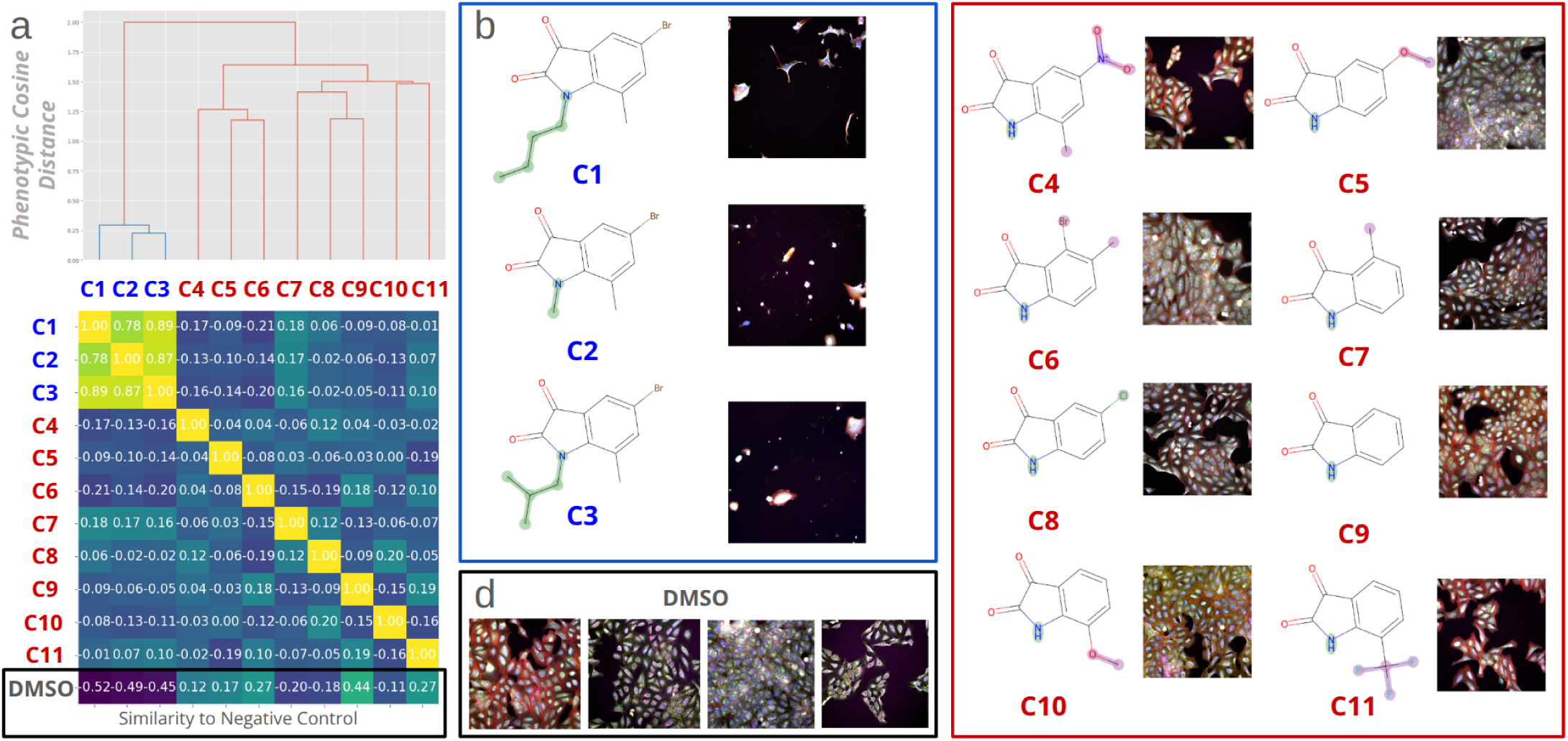
**(a)** *Example of Phenotypic Hierarchical Clustering of Compounds with the same Murcko Scaffold.* The distance matrix indicates the cosine similarities between phenotypes induced by these compounds and both the dendrogram and the matrix highlight two main phenotypic clusters: the first one is a group of very similar phenotypes while the second displays a variety of sub-phenotypes as highlighted by the matrix diagonal. The last row of the matrix indicates the distances to the negative control (DMSO). **(b)** Further investigation of the images and the structures shows that the first cluster includes 3 apparently toxic compounds each featuring an aliphatic chain highlighted in green. **(c)** The second cluster presents *compounds that induce different phenotypes.* Highlighted in green is the common protonated nitrogen atom. In purple the different substitution that could explain the different phenotypes induced. **(d)** Examples of negative control with DMSO, a mock treatment.

### Compounds reproducing the positive control phenotype tend to hit targets in the same pathway

We further investigated if compound perturbations reproducing the phenotype of a positive control were modulating the same biological pathways. A biological or signaling pathway is a series of biomolecular interactions, such as biochemical reactions or signaling events, through which cells respond to internal or external stimuli. Pathways regulate critical cellular processes including metabolism, gene expression, and cell signaling, enabling coordinated cellular behavior. To illustrate this hypothesis, we considered data from a fluorescence cell-based assay screen to identify inhibitors of the EGFR pathway (CHEMBL1613808). We chose Dezmapinod, a non-specific inhibitor of mitogen-activated protein kinases (MAPK), as our positive control (**Figure 6a**). Then, we considered the top 5% compounds (15 compounds out of 300 that appeared both in CHEMBL1613808 and JUMP-CP) with highest phenotypic similarity to Dezmapinod. Among these 15 compounds (**Figure 6b**), three were known to specifically inhibit proteins within the same pathway: EGFR for AG1478, BRD4 for Sb-202190, and MAPK8 for SP600125 (**Figure 6c**). We explored a prior-knowledge database to connect these genes into a signaling pathway. Among all the existing databases, we focused on SIGNOR ^59^ (see <u>Methods</u>). Although BRD4 was not part of the initial SIGNOR signaling pathway, it is known to modulate MYC activity and conversely ^60,61^ and therefore added to the network. These findings on a specific example suggest that compounds targeting the same pathway, even when they act on different proteins, can produce a similar phenotype.

**Figure 6:**
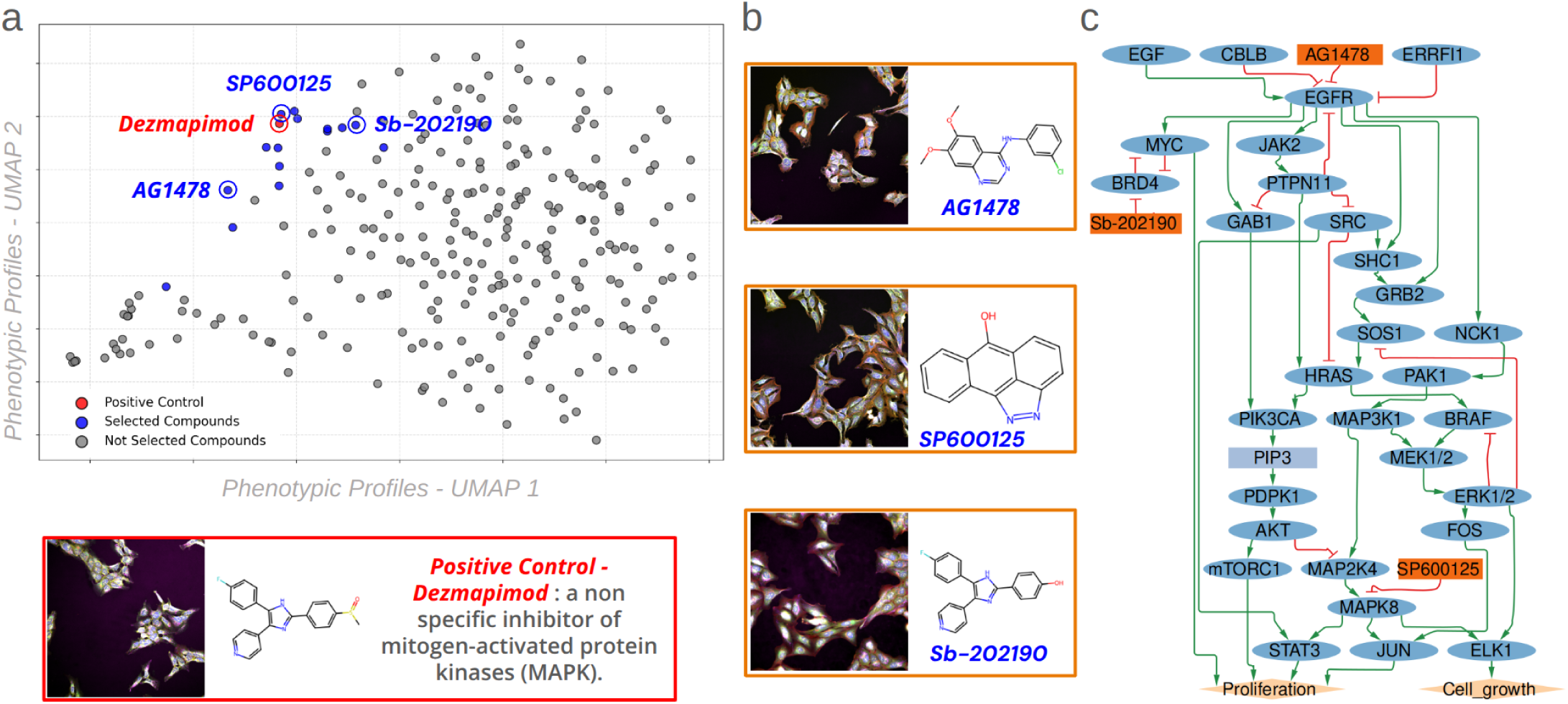
Example of a screen where compounds reproducing the positive control phenotype hit different targets of the EGFR pathway. **(a)** UMAP^54^ of phenotypic profiles of compounds from the CHEMBL1613808 screen, designed to identify *in cellulo* inhibitors of the EGFR pathway. Positive control (***Dezmapimod***) is shown in red, while compounds selected by our method are shown in blue. **(b)** Examples of selected compounds (SC) include known EGFR pathway inhibitors, acting through distinct mechanisms of action and targeting different genes involved in the pathway. **(c)** The EGFR signaling pathway extracted from the SIGNOR database. Targets of the selected compounds are highlighted in blue, illustrating their diverse role within the pathway and confirming why structural similarity between compounds is not always expected.

### Compounds targeting the same pathway tend to induce related phenotypes

We then aimed to evaluate the extent to which the previous observations, made on a single pathway and given a specific screen positive control, generalizes to any signaling pathways. To this end, we cross-referenced genes associated with the 30 pathways from MSigBD that had more than 10 known compound–gene interactions in BindingDB targeted by compounds found in JUMP-CP. For each pathway, we identified a few hundred compounds targeting up to a few tens of genes (see <u>Methods</u>). We then compared the phenotypic similarities between those compounds against the phenotypic similarity between random compounds in JUMP-CP for each pathway (**Figure 7a**). As a result, we observed that compounds targeting the same pathway exhibit a significantly higher frequency of extreme phenotypic similarities, both highly similar and almost opposite, compared to random pairs. This result is consistent across every one of the 30 studied pathways (see <u>Methods</u>, **Supplementary Fig. 19** and **Table 1** for all distributions and statistical details). On one hand, these findings confirmed the hypothesis that compounds targeting the same pathway could be found through their phenotypic similarity. On the other hand, interestingly, it showed that a high dissimilarity could also be used as a criteria for selection. One explanation for the latter is that compounds targeting the same pathway may trigger opposite downstream effects, leading to opposite cellular responses because of the role of the target on the pathway. Illustrating it on the G2M pathway (**Figure 7**), the previously described procedure led to a significantly high amount of compounds inducing both highly similar or highly dissimilar phenotypes (**Figure 7a**). We then examined the effect of four well-characterized compounds known to interact with specific targets in this pathway and present in the JUMP-CP dataset (SCHEMBL1578316, Reversine, NSC-625987, and Gnf-5; **Figure 7b**) that are displaying both similar and opposite phenotypic profiles (in terms of cosine similarities, see **Figure 7c**). To this end, we constructed the network interaction with the 21 G2M pathway targeted genes using OmniPath interactions via the NeKo tool^61^ and considered the subnetwork with only genes linked to those compounds (**Figure 7d**). We found that the direction of their phenotypic similarities, either similar or opposite phenotypic profiles (**Figure 7c**), is coherent with their downstream effects on the interaction network. For instance, Reversine inhibits an inhibitor (AURKB) of cell proliferation (BIRK5), leading to a pro-proliferative outcome, whereas NSC-625987 inhibits CDK4, blocking proliferation. These two compounds target the same pathway with opposite effects. Even if these observations cannot, yet be generalised to any pair of compounds, the visualization of their effect on networks highlights which mechanisms each of these compounds targets and can explain the observed phenotypes. The presence of this bimodal effect (highly similar / highly opposite phenotypic profiles)^26^ in our representation of compounds displays the power and the complexity of biological information captured by phenotypic profiles.

**Figure 7:**
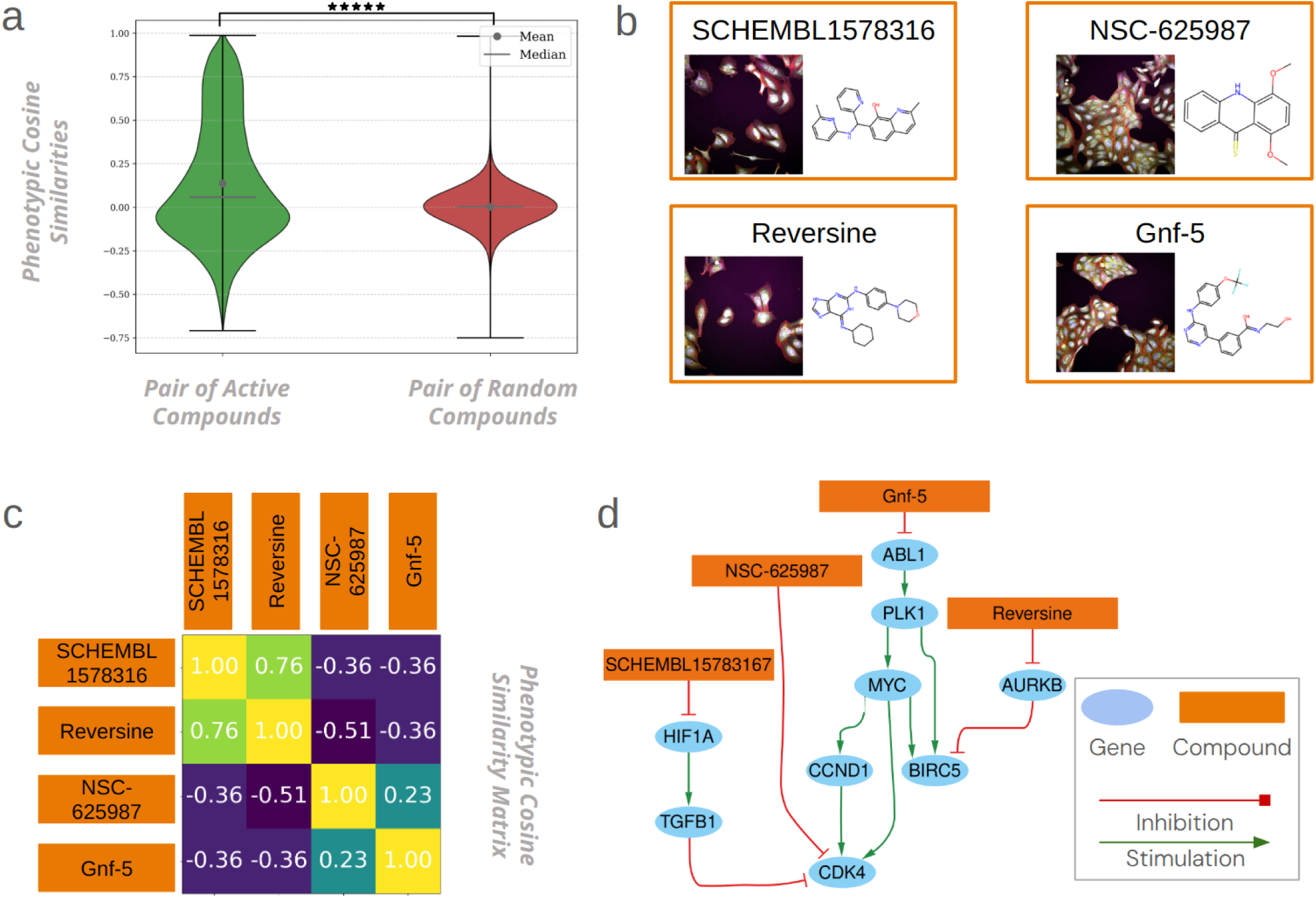
Phenotypic similarities and dissimilarities of compounds targeting G2M pathway genes. **(a)** Violin plot of phenotypic similarity distributions: in blue, similarities between all pairs within the 530 compounds active against the G2M pathway; in orange, similarities between compound pairs inactive against the G2M pathway; in green, pairs consisting of one active and one inactive compound. **(b)** Cellular image (Phenotypes) and chemical structures of the four compounds. **(c)** Cosine similarity matrix showing the phenotypic similarity among the four compounds, each targeting different G2M pathway genes. Negative signs show opposite phenotypic profiles whereas positive signs show similar phenotypic profiles **(d)** Network of gene–gene interactions among G2M pathway genes targeted by JUMP-CP compounds. G2M pathway genes were obtained from MSigDB, interactions from OmniPath, and the network was constructed using NeKo. Red edges indicate inhibition, green edges indicate stimulation, and purple edges a bimodal effect.

**Table 1:**
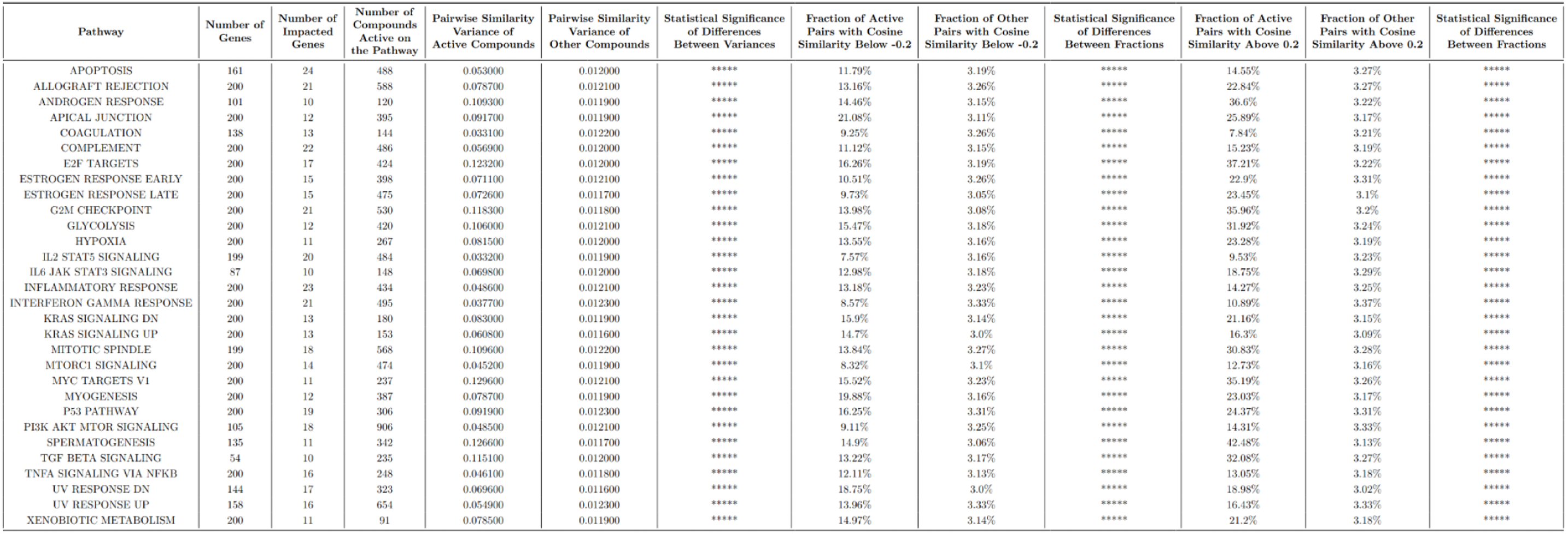
Statistical Comparison of Phenotypic Similarities Between Pathway-Targeting and Random Compound Pairs.

## Discussion

QSAR-based library selection methods rely on the premise that chemically similar compounds often share biological activities. While valuable, this approach inherently limits the chemical diversity of selected compounds to those resembling the reference compound in structure. Through an extensive evaluation of deep image encoding and data processing methods, we identified an approach that enables robust pairwise comparison of phenotypic profiles of compounds across the entire JUMP-CP dataset while preserving biological signals and mitigating batch effects. Leveraging this approach, we were able, for the first time, to harness this vast dataset to develop a training-free transfer learning approach for the efficient selection of bioactive compounds for screening assay. Our method was extensively validated across 65 diverse high-throughput assays from both internal and public databases, covering *in cellulo* and *in vitro* bioactivities, underscoring the broad applicability and robustness of Cell Painting phenotypic profiling across biological systems. Importantly, because it is training-free, this approach is straightforward to use, as demonstrated by our web-based tool Phenoseeker, which allows researchers to simply submit the name of a positive control from their drug discovery project and retrieve, with a single click, a list of JUMP-CP compounds most likely to exhibit similar bioactivity.

Interestingly, we enabled, for the first time, a structure-activity relationship analysis spanning the largest publicly available dataset. On one hand, we systematically identified activity cliffs, where slight modifications in compound structure result in significant phenotypic variations, highlighting crucial compound properties that would remain undetected by structure-based approaches alone. By doing so, we demonstrated that deep phenotypic profiles can provide subtle yet critical information at the atomic level. On the other hand, we found that compounds with entirely different structures can target genes in the same pathway, inducing either significantly similar or distinct phenotypic profiles. This was validated across 30 pathways, hundreds of genes, and thousands of compounds. We illustrated this through examples leveraging known compound-to-gene and gene-to-gene interaction networks, reinforcing why phenotypic similarity and dissimilarity represent meaningful metrics for selecting bioactive compounds with a diverse range of molecular structures.

While we demonstrated the effectiveness of the proposed approach, some limitations should be acknowledged. First, although JUMP-CP is the largest publicly available dataset of its kind, it provides a phenotypic view based on a single cell line: U2OS. This constraint means that compounds targeting genes not expressed in U2OS may not produce the relevant phenotypic profiles, potentially leading to missed selections. Another limitation is that compound selection is restricted to the 112.480 compounds in the JUMP-CP library. While substantial, this represents only a small subset of available compounds and an even smaller fraction of the theoretical or even the smaller drug-like chemical space. A possible solution to this limitation would be to train a cross-modal model, which could predict a phenotypic profile for any given compound structure, allowing us to identify similar JUMP-CP phenotypes even for novel compounds. We recently trained such a model based on CLIP on the full JUMP-CP dataset^52^, but it has not yet been extensively evaluated as a compound selection tool in existing screens. Moreover, learning the complex relationship between chemical structure and cell phenotype remains challenging due to the discontinuities introduced by activity cliffs and plateaus we observe in this work. While such a cross-modal model could estimate the phenotype of a given structure, it would not generate new structures likely to induce a desired phenotype—an ultimate goal in drug discovery. To achieve this, a generative model should be conditioned on target phenotypic profiles to design compounds with targeted bioactivity, and the approach we proposed here could provide a valuable foundation for such training.

Our findings underscore the power and scalability of the proposed compound pre-selection approach in both academic research and drug discovery. By harnessing phenotypic response as a biological proxy of compounds bioactivities, our method transcends the constraints of structural similarity, venturing into unexplored regions of chemical space to uncover novel bioactive compounds. This leap beyond traditional QSAR methods not only enhances structural diversity but also reveals drug-like candidates that might otherwise remain hidden. While structure-based approaches rely on chemical resemblance to predict bioactivity, inherently restricting exploration, our method captures diverse biological mechanisms and drug-target interactions, offering a powerful and complementary alternative to conventional selection strategies.

## Methods

### Cell Painting dataset

**Cell Painting** is a multiplexed high-content imaging assay designed to capture comprehensive phenotypic signatures of cellular responses. This method employs fluorescent dyes targeting multiple cellular compartments—including nuclei, cytoskeleton, mitochondria, endoplasmic reticulum, and Golgi apparatus.

We used the JUMP-CP dataset from the eponymous consortium ^38^, which encompasses more than 136,000 chemical and genetic perturbations represented by over 5 million images acquired across 12 laboratories. These experiments were performed following the standardized Cell Painting assay protocol described in Bray et al ^5^ Each Cell Painting image consists of five grayscale channels, each highlighting one or several key cellular organelles. Each assay plate includes negative controls (DMSO mock treatment) with 32 replicates on 384-well plates and 128 replicates on 1536-well plates, and positive controls comprising eight selected compounds (**Supplementary Table 1**), each with 4 replicates on 384-well plates and 16 replicates on 1536-well plates.

Specifically, we use the compressed version from Watkinson et al ^52^. It encompasses 112.480 unique chemical compounds.

### Raw feature extraction from images

We utilized three main categories of models to extract features from the 5-channel Cell Painting images:

#### (1) Handcrafted feature extraction

We used **CellProfiler** ^42^, a software designed to extract handcrafted morphological features at the single-cell level. After segmenting individual cells, features were computed per cell and then aggregated at the well level. For this study, we relied on precomputed and normalized features made publicly available by the Broad Institute (https://broadinstitute.github.io/jump_hub/howto/11_retrieve_profiles.html).

#### (2) Deep learning models pretrained on natural images

Two neural network architectures originally trained on natural image datasets were adapted to our Cell Painting dataset. To accommodate the five-channel format of Cell Painting, the first convolutional layer’s weights initially trained on the RGB channels were duplicated: channels 4 and 5 used copies of the weights from the red (R) and green (G) channels, respectively. Note that the specific choice of the order of Cell Painting channels does not affect the model performance, as long as consistency is maintained during inference.

- **ResNet50** ^43^, a convolutional deep neural network architecture that leverages residual connections widely used to extract features with robust performance across diverse tasks.
- **DINOv2** ^44^, a state-of-the-art self-supervised Vision Transformer (ViT) that employs a teacher-student framework to learn robust, transferable representations from unlabeled data, thereby enabling efficient feature extraction across a range of tasks. We evaluated both the Small (DINOv2-S, 21M parameters) and the Giant (DINOv2-G, 1300M parameters) versions of DINOv2 to assess the impact of model depth and complexity on feature representation.

#### (3) Deep learning models pretrained on microscopy images

We also explored two architectures specifically trained on microscopy data, offering potential advantages due to domain-specific pretraining:

- **ChAda-ViT (Channel Adaptive Vision Transformer)**^45^, a ViT model tailored for microscopy images with heterogeneous input channels (ranging from 1 to 10). ChAdA-ViT employs cross-channel attention mechanisms to adaptively learn joint representations, making it well-suited to multi-channel assays like Cell Painting.
- **OpenPhenom** ^46^, an open-source Masked Autoencoder (MAE)^63^ architecture explicitly trained on Cell Painting images. Developed using millions of Cell Painting images, OpenPhenom learns robust and generalizable representations through a self-supervised reconstruction task.

A detailed summary of the compound selection results for all models tested is provided in the supplementary material (**Supplementary Fig. 16**).

Several notable observations emerged from our analysis. Although CellProfiler shows excellent performance for retrieving replicates of identical perturbations and effectively reducing batch effects, this may partly result from differences in normalization procedures: CellProfiler features provided by the Broad Institute were normalized using all available negative controls from the original JUMP-CP dataset, whereas our analysis relied on approximately 70% of the wells and 6 instead of 9 images per well, as we used the compressed JUMP-CP dataset version published by Watkinson et al. Interestingly, neural networks pre-trained on natural images show strong performance, despite never having encountered Cell Painting data during their training. Among these, DINOv2 stands out by producing highly discriminative phenotypic features. ChAda-ViT achieves good results, considering its relatively small training set of ∼100k microscopy images (including around 55k fluorescence microscopy images). Overall, DINOv2-Giant appears to be the optimal model for our selection method, slightly outperforming OpenPhenom.

### Features alignment on DMSO

We evaluated several feature-wise alignment methods to standardize phenotypic profiles, aiming to minimize batch effects while preserving biological signals. The tested methods included:

**Rescaling (Res)**: Each feature *x* (*i* ∈ [|1, *n_features_*|], *n_features_* being 192, 384, 1515 or 2048 depending on the encoder) is linearly scaled to a defined range, either [0, 1] or [-1, 1], using:

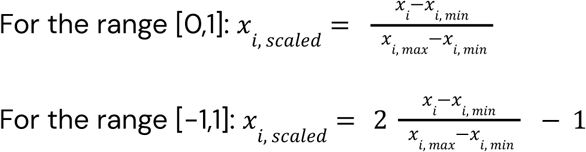

The minimum and maximum across all samples.

#### Removal (Rem)

Features with low variability across samples were removed, based either on standard deviation (std) across samples, or mean absolute deviation (MAD) across samples, using thresholds.

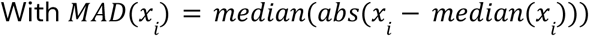

#### Z-score normalization (Z)

Each feature *x* is standardized to zero mean and unit variance using its mean across samples (µ) and standard deviation across samples (σ):

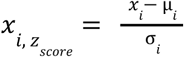

#### Robust Z-score normalization (rZ)

A robust variant using median across samples and median absolute deviation across samples, less sensitive to outliers:

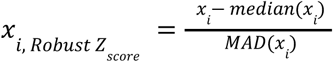

#### Inverse normal transform (Int)

Features are mapped onto a standard normal distribution by applying the rank-based inverse normal transformation:

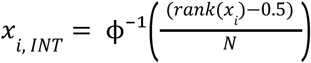

where ϕ⁻¹ is the inverse cumulative normal distribution and *N* is the number of samples.

#### Sphering (Sph)

We whiten each plate’s feature matrix *X* ∈ *R^*n*×*d*^* (with *n* samples, *d* features) by centering and then multiplying by the inverse square root of its covariance. Concretely:

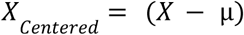

where µ is the feature-wise mean vector.

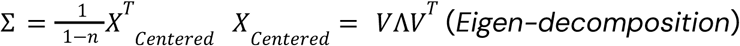

where Σ is the covariance matrix and Λ = *diag*(λ_1_, …, λ*_d_*)

Two whitening methods were tried :

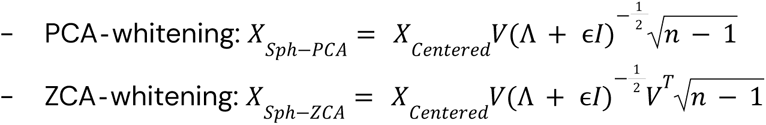

Where ϵ = 10^−6^ is a small “fudge factor” to avoid division by zero on near-rank-deficient plates.

ZCA preserves each sample’s orientation in the original feature space, while PCA rotates onto the principal-component axes.

Each method was evaluated individually as well as in various combinations. For Z-score and Sphering methods, transformations were fitted either solely on negative controls (DMSO wells) or across all wells on a given plate. All normalization procedures were performed on a plate-wise and feature-wise basis.

### Metric used to compare phenotypic profiles

In this article, we used cosine similarity to compare phenotypic profiles due to its ability to identify biologically related samples based on the similarity in their patterns of change rather than on the magnitude of these changes^65^. Additionally, cosine similarity provides directional information, capturing the relationship orientation between two phenotypic profiles.

The cosine similarity is defined as :

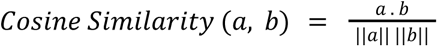

Where *a*. *b* is the scalar product between *a* and *b*, and ||. || is the Euclidean norm of the vector.

### Metrics and data used to compare profiling methods

We employed specific metrics tailored to our downstream application—bioactive compound selection through ranking and retrieval. Although these metrics are not flawless, they effectively capture our primary objectives. In particular, we utilized the **mean average precision (mAP)** ^64^ metric, a standard evaluation measure widely employed in information retrieval and ranking tasks. Specifically, the average precision (AP) for a given query is calculated by averaging the precision values at each position in the ranked list where a relevant item appears. The mAP is then obtained by averaging AP scores across all queries, effectively capturing both the quality of retrieved items and their ranking order. Higher mAP scores thus indicate superior retrieval performance and accuracy.

For a given query (a well phenotypic profiles), the Average Precision (AP) is defined as:

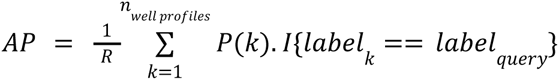

Where :

- *P*(*k*) is the precision at rank *k* (here,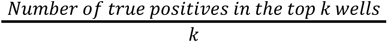)
- *R* the total number of relevant (ie, with the same label as the query one) well profiles
- *I*{.} the indicator function (1 if condition holds, 0 otherwise)

To quantify batch effect reduction, mAP was computed using phenotypic profiles from both negative (DMSO) and positive control wells, with experimental batch identifiers serving as labels. Conversely, to assess the preservation of biological signals, mAP was computed exclusively on positive control wells, using compound identities as labels. The mAP was not computed with sample wells to avoid potential bias due to sample phenotypic profiles distributions.

We can observe that all 8 positive controls do not have the same phenotypic impact on cells (**Supplementary Fig. 1**, **Supplementary Fig. 2**). Indeed, some have strong effects (JCP2022_046054, JCP2022_037716 or JCP2022_035095), some have weaker effects (JCP2022_012818 or JCP2022_064022) and some are not distinguishable from DMSO negative control (JCP2022_085227, JCP2022_050797). This explains the huge difference in mAP value between those compounds and the average mAP across those 8 compounds being only around 0.5 for best models and normalisations. The strong batch effect observed on **Supplementary Fig. 2** also explains mAPs values.

For computational efficiency, we initially evaluated all possible combinations of normalization methods (over 5,000 in totals) using a subset of five plates per laboratory, each selected from distinct experimental batches. Subsequently, the 100 best-performing combinations were re-evaluated using approximately half of the JUMP-CP dataset (∼600 plates), explicitly chosen to maximize stratification across experimental batches and laboratories. This evaluation process was conducted independently for each feature extractor: ChAda-ViT, ResNet50, DINOv2-G, DINOv2-S, and OpenPhenom.

Figure 1 shows the performance results for all tested combinations, while **Supplementary Fig. 3** presents detailed results for the 100 best-performing combinations evaluated on the larger subset (∼600 plates). Importantly, results remained consistent from the initial smaller set of plates (45 plates) to the expanded validation (600 plates), supporting the robustness of our normalization approach.

### Compound structural representation using Morgan fingerprints

To describe the structural topology of chemical compounds, we computed Morgan fingerprints^56^ using the RDKit toolkit. Specifically, Morgan fingerprints were generated with a radius of 2 and a bit-vector length of 1024. This fingerprinting method encodes structural information by iteratively considering atom neighborhoods within the specified radius, capturing both local chemical environments and global topological properties of the compound. The similarity between compounds was quantified using the Tanimoto similarity^57^, a widely adopted measure for comparing molecular fingerprints. The Tanimoto similarity assesses the overlap of structural features between pairs of compounds.

It is defined as:

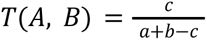

where:

- *a* and *b* are the number of bits set to 1 in the fingerprints of compounds *A* and *B*, respectively,
- *c* is the number of bits set to 1 in both fingerprints.

### Chemical diversity of selected compounds

To compare the distribution of Tanimoto similarities in **Figure 3**, we first checked normality of the paired differences using the Shapiro–Wilk test at a significance level of α=0.05. Because most differences did not follow a normal distribution (p<0.05, see **Supplementary Table 2**), we employed the nonparametric Wilcoxon signed-rank test with a one-sided alternative hypothesis (e.g., “Top 5% Structures” > “All Tested Compounds”). This approach provides a robust evaluation of whether one distribution exhibits systematically higher (or lower) Tanimoto similarity compared to another.

The resulting *p*-values were then converted into significance thresholds (*p* < 0.05, < 0.01, < 0.001, etc.), which are visually depicted in box plots using the star notation (e.g. ∗, ∗∗, ∗∗∗, etc.). All test values and raw p-values are displayed in **Supplementary Table 2**

### Chemical diversity of JUMP-CP

We compute a UMAP projection of the Morgan fingerprint of all compounds in JUMP-CP alongside those from ChEMBL and DrugBank to assess the extent of the chemical space covered by JUMP-CP compounds relative to known or previously studied compounds (see **Supplementary Fig. 18**). These UMAP projections show that JUMP-CP compounds broadly overlap with both ChEMBL and DrugBank while also occupying distinct regions in chemical space, underscoring the shared and novel structural diversity within the JUMP-CP dataset.

### Assays description

#### Screen

A high-throughput experimental procedure to rapidly evaluate the activity of a library of compounds.

#### Assay

A standardized test that quantifies or qualifies the interaction, activity, or presence of a compound or biomolecule, used to validate screening results.

##### Institut Curie BioPhoenix screens (Supplementary Fig. 5)

We included assays from the Institut Curie’s BioPhoenix screening platform based on specific criteria. Assays were selected if at least 100 compounds from the JUMP-CP dataset were screened, and if at least five active compounds (hits) were identified in the original assay. These screens were all conducted using cellular models, primarily employing imaging-based readouts, although some relied on fluorescence-based methods such as CellTiter-Glo. Many assays featured overlapping sets of compounds, typically originating from shared chemical libraries like the Prestwick library. Despite this overlap, hits identified varied significantly from one assay to another, even when the same compounds were tested. In total, 16 BioPhoenix screens met these criteria.

##### ChEMBL screens (https://www.ebi.ac.uk/chembl/) (Supplementary Fig. 6)

We also incorporated assays from the ChEMBL database, applying the same criteria as those used for BioPhoenix assays. Assays included had at least 5 active and 100 inactive compounds also present in the JUMP-CP dataset. To focus specifically on challenging screening tasks, we retained only assays with a hit rate lower than 35%. The selected ChEMBL assays represent diverse experimental methodologies and biological contexts, covering *in vitro* and *in cellulo* systems. Overall, this resulted in 44 selected ChEMBL screens.

##### Lit-PCBA benchmark ^55^ (Supplementary Fig. 7)

Finally, we utilized the Lit-PCBA dataset for additional benchmarking. From Lit-PCBA, we retained only targets with at least 10 active compounds and a minimum of 100 total compounds present in the JUMP-CP dataset. This filtering resulted in five targets meeting our criteria: ALDH1, MAPK1, PKM2, CDR, and ESR1-antagonist.

Collectively, Lit-PCBA target, ChEMBL and BioPhoenix assays encompassed significant chemical diversity, covering more than 10,000 unique compounds from the JUMP-CP dataset—representing around 10% of the dataset in its entirety.

A supplementary table for each source (BioPhoenix, ChEMBL, Lit-PCBA) provides detailed information including the screen or target identifiers, assay type, hit rate, and total number of tested compounds.

Additionally, **Supplementary Fig. 13** illustrates the proportion of the JUMP-CP dataset evaluated across all selected assays.

### Normalized enrichment factor

To evaluate the ability of our method to select bioactive compounds, we compute enrichment factors ^65^. We chose a threshold of 5% to select at least 5 compounds for all screens where we have 100 tested compounds (but results are consistent across thresholds, see **Supplementary Fig. 4**). Those values were normalized to help the comparison across screens with a very diverse number of compounds and hit rate. A normalized enrichment factor (nEF) of 100% means the best possible selection of compounds.

The enrichment factor at 5% is computed as:

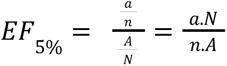

where:

- *a* is the number of active compounds in the top 5% selected compounds,
- *n* is the total number of compounds selected (5% of *N*),
- *A* is the total number of actives in the dataset,
- *N* is the total number of compounds tested.

The Normalized Enrichment Factor (nEF) is then defined as:

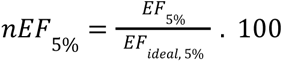

where: *EF*_*ideal*, 5%_ is the theoretical maximum enrichment at 5% (i.e., when all selected compounds are active or all active compounds are selected).

### Bemis-Murcko scaffolds and phenotypic subclusters

To systematically group compounds according to their structural similarity, we computed molecular scaffolds using RDKit’s implementation of the Bemis–Murcko scaffold definition. Note that RDKit’s implementation differs slightly from the original Bemis–Murcko definition^58^: it retains the first atom of exocyclic substituents attached via double bonds, distinguishing certain structural motifs (e.g., differentiating between C1CC1=O and C1CC1). Further details about these differences are documented in the RDKit community discussion (https://github.com/rdkit/rdkit/discussions/6844).

Once scaffolds were generated within each scaffold-defined group, we performed hierarchical clustering to further subdivide compounds into distinct phenotypic subgroups. We retained groups where each cluster contained at least three compounds to ensure sufficient representation for meaningful analysis. To explore the robustness of our approach, we systematically tested scenarios with two, three, four, or five clusters per scaffold, resulting in corresponding minimal scaffold group sizes of 6, 9, 12, or 15 compounds.

Detailed statistical results from these different clustering configurations are provided in **Supplementary Fig. 11**.

Scaffolds were selected as exposing subclusters with substantially different phenotypes when intra-cosine phenotypic similarity was higher than inter-cosine phenotypic similarity by at least 0.2 (in **Figure 4**). The value 0.2 corresponds to a cosine similarity greater than 3 standard deviations from the mean of the whole dataset (see **Supplementary Fig. 15** for distribution of similarity across all JUMP-CP compounds).

### Pathways and networks construction

The EGFR pathway used in **Figure 6** corresponds to the signaling network provided by The SIGnaling Network Open Resource (SIGNOR 3.0, https://signor.uniroma2.it) with a confidence superior to 0.2 for all interactions. Additionally, the Brd4 gene was manually included based on evidence from previous literature facts ^60,61^.

For **Figure 7** and **Table 1**, gene sets were obtained from The Molecular Signatures Database (MSigDB, https://www.gsea-msigdb.org/), a curated collection of annotated gene sets derived from diverse biological sources, including published experimental data, expert-curated pathways, and computational predictions. We used the Hallmarks gene sets that focus on cancer but it can be generalized. Compound-target relationships were retrieved from BindingDB (https://www.bindingdb.org/), a comprehensive database of experimentally validated compound–gene interactions. We retained only compounds present in both BindingDB and the JUMP-CP dataset, along with genes targeted by these remaining compounds. Detailed statistics regarding genes, filtered genes, and remaining compounds are presented in **Table 1**.

The gene interaction network shown in **Figure 7** was constructed using the NeKo tool^62^, which aims at reconstructing connected networks from curated prior-knowledge databases (i.e., OmniPath (https://omnipathdb.org/), a comprehensive resource for intra- and intercellular signaling knowledge). We built the network specifically with the 21 genes from the G2M pathway targeted by JUMP-CP compounds and focused on the SIGNOR database only. This complete NeKo-generated network is available in **Supplementary Fig. 17** The subnetwork in **Figure 7**. highlights interactions relevant to the four selected compounds.

### Pathways & pPhenotypes table and statistics

For each gene set corresponding to a given biological pathway, we retained only those pathways in which at least 10 distinct genes were targeted by at least 50 compounds from the JUMP-CP dataset, resulting in 30 pathways for further analysis. For each selected pathway, we computed pairwise cosine similarities between phenotypic profiles of all compounds known to target genes within that pathway. As a comparative control, we also computed cosine similarities among a random set of non-targeting compounds, sampled at a ratio of 10 non-targeting compounds per active compound. The resulting distributions of cosine similarities for each pathway, including both active and random compound pairs, are shown in **Supplementary Fig. 19**.

To determine whether active compounds exhibited more extreme similarity or dissimilarity, compared to random compound pairs, we first tested differences in variance between these two distributions using Levene’s test. Additionally, to assess whether active compound pairs were significantly enriched in highly similar (cosine similarity > mean + 3 standard deviations, see **Supplementary Fig. 15** for distribution of similarity across all JUMP-CP compounds) or highly dissimilar pairs (cosine similarity < mean - 3 standard deviations, see **Supplementary Fig. 15** for distribution of similarity across all JUMP-CP compounds), we calculated the percentage of pairs meeting these thresholds in both active and random distributions, and assessed statistical significance using Fisher’s exact test. Detailed statistical results, including test statistics and p-values, are provided in **Table 1**.

## Data availability

JUMP-CP dataset is publicly available (see: https://broadinstitute.github.io/jump_hub/). Screens data from ChEMBL^66^ is publicly available; we used version 35 of this database (see: https://www.ebi.ac.uk/chembl/). Screens data from BioPhoenix are not publicly available. Compounds-targets interactions are publicly available (see: https://www.bindingdb.org/). The gene sets are publicly available (see: https://www.gsea-msigdb.org/).

## Code availability

We provide scripts at https://github.com/mxfly14/2025_sanchez_phenoseeker for the automated preprocessing of the data, test and evaluation of normalization methods and reproduction of the figures and plots. The code to download the compressed version of JUMP-CP is available at https://github.com/gwatkinson/jump_download.

## Acknowledgements

This work has received support under the program « Investissements d’Avenir » launched by the French Government and implemented by the ANR, with the references: ANR-10-LABX-54 MEMO LIFE ANR-11-IDEX-0001-02 PSL* Research University, MS was funded by Iktos and ANRT under the CIFRE program. This work was granted access to the HPC resources of IDRIS under the allocation 2020-AD011011495 made by GENCI. We thank Gabriel Watkinson for helping with the JUMP-CP dataset.

## Author Contributions

MS designed the computational framework and performed all the analyses. NB, IB, EC and IS helped for the analyses. HT and GB contributed to the discussion. AG and LC conceived the project and supervised the work. MS and AG wrote the manuscript. All authors revised the manuscript.

