## Supplementary Information for "Large Scale Cell Painting Guided Compound Selection Reveals Activity Cliffs and Functional Relationships"

#### Supplementary Figures

JCP2022_085227 :

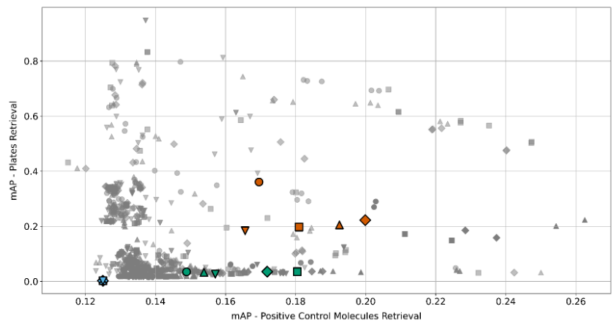


JCP2022_064022 :

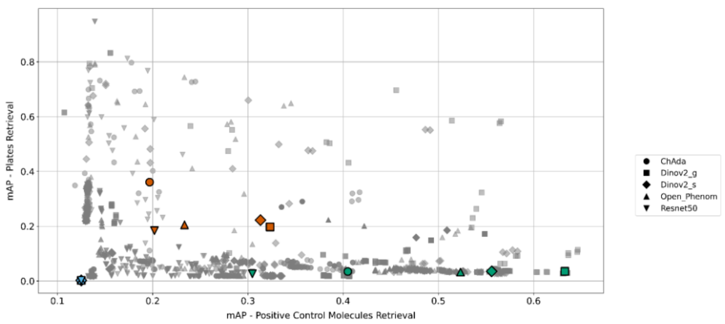


JCP2022_050797 :

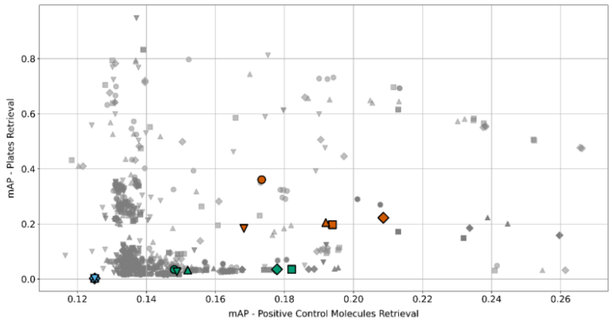


JCP2022_046054 :

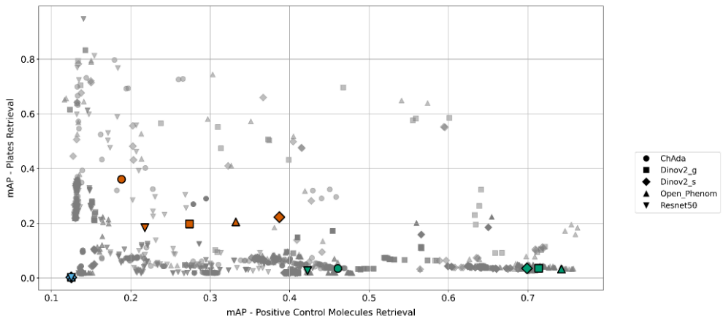


JCP2022_037716 :

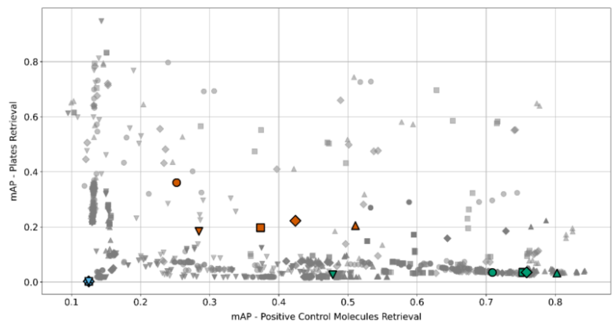


JCP2022_035095 :

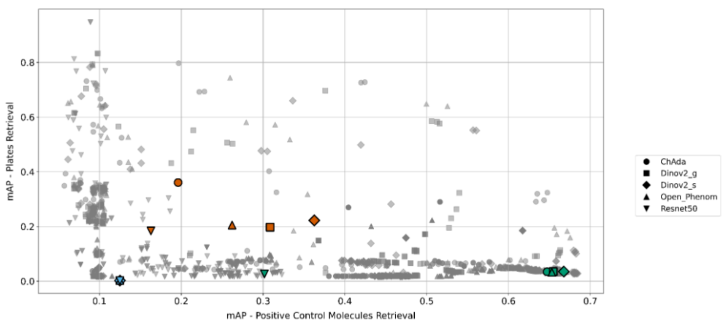


JCP2022_012818 :

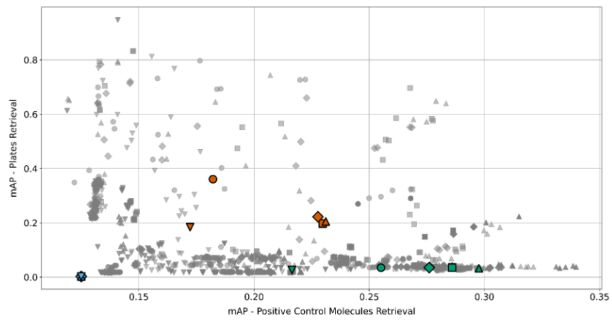


JCP2022_025848 :

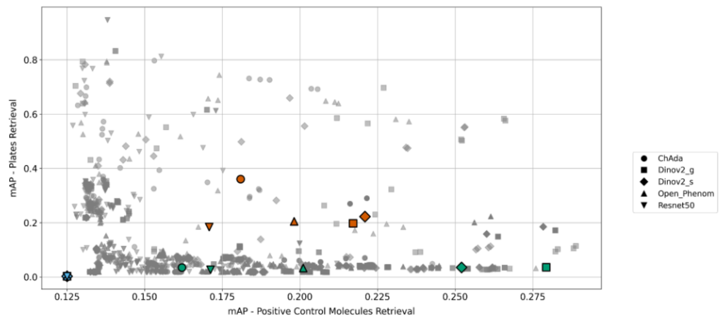

**Supplementary figure 1:** mAPs values for all 8 positive control compounds present in the 11 JUMP sources. mAPs for plate retrieval are always the same, but mAPs for positive control compounds retrieval are individual.

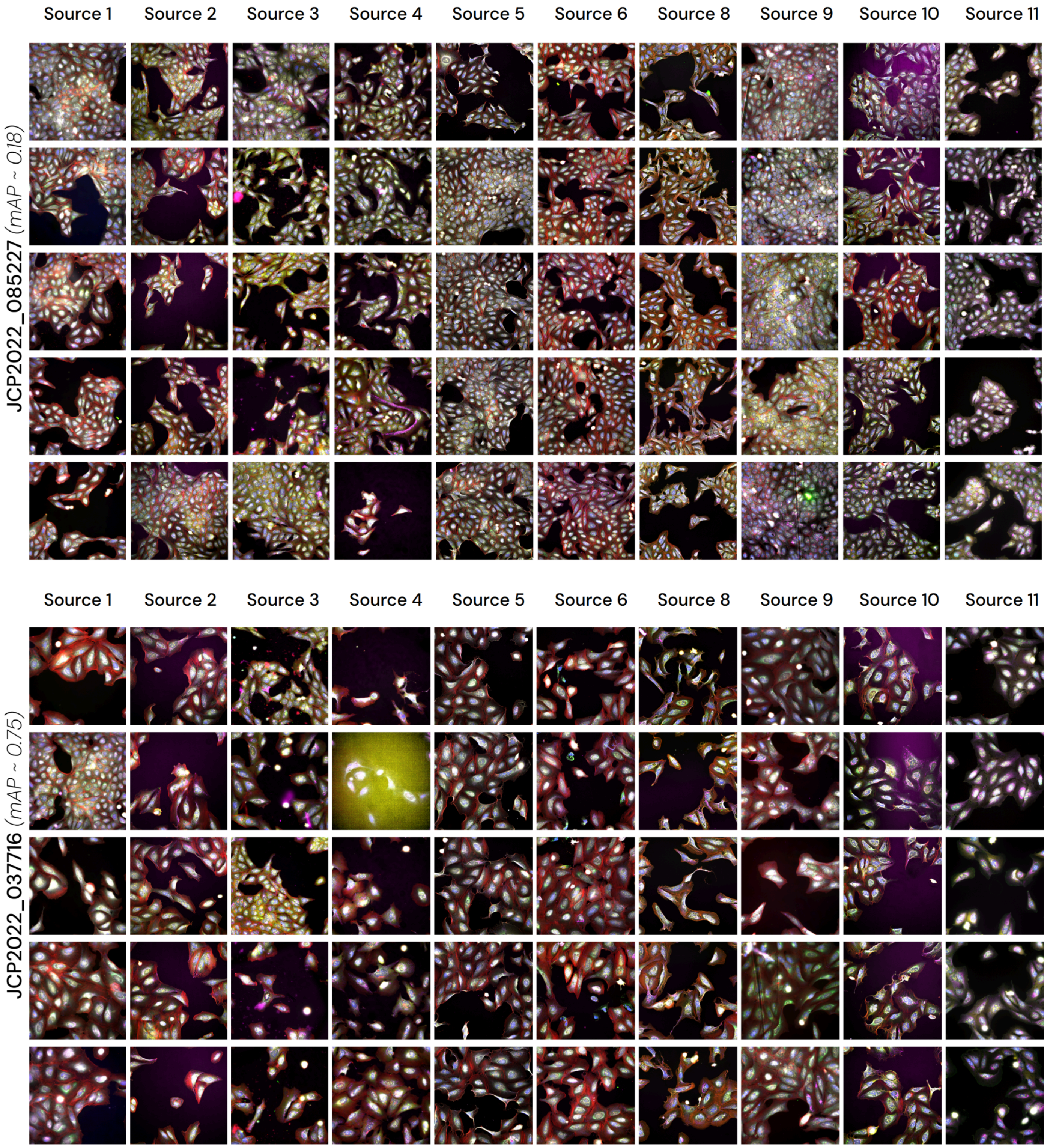

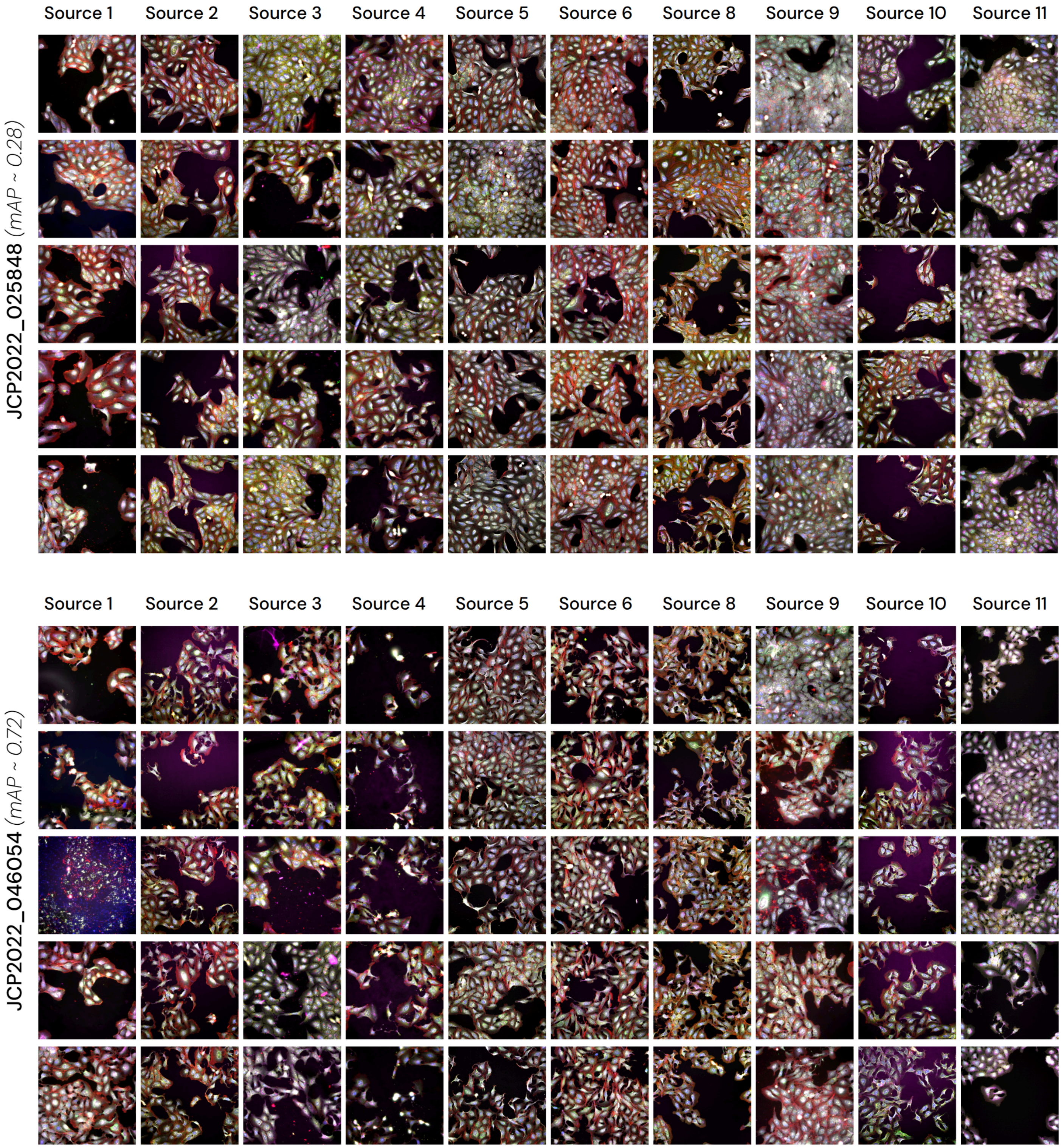
Source 1
Source 2
Source 3
Source 4
Source 5
Source 6
Source 8
Source 9
Source 10
Source 11
JCP2022_025848 (mAP ~ 0.28)
Source 1
Source 2
Source 3
Source 4
Source 5
Source 6
Source 8
Source 9
Source 10
Source 11
JCP2022_046054 (mAP ~ 0.72)

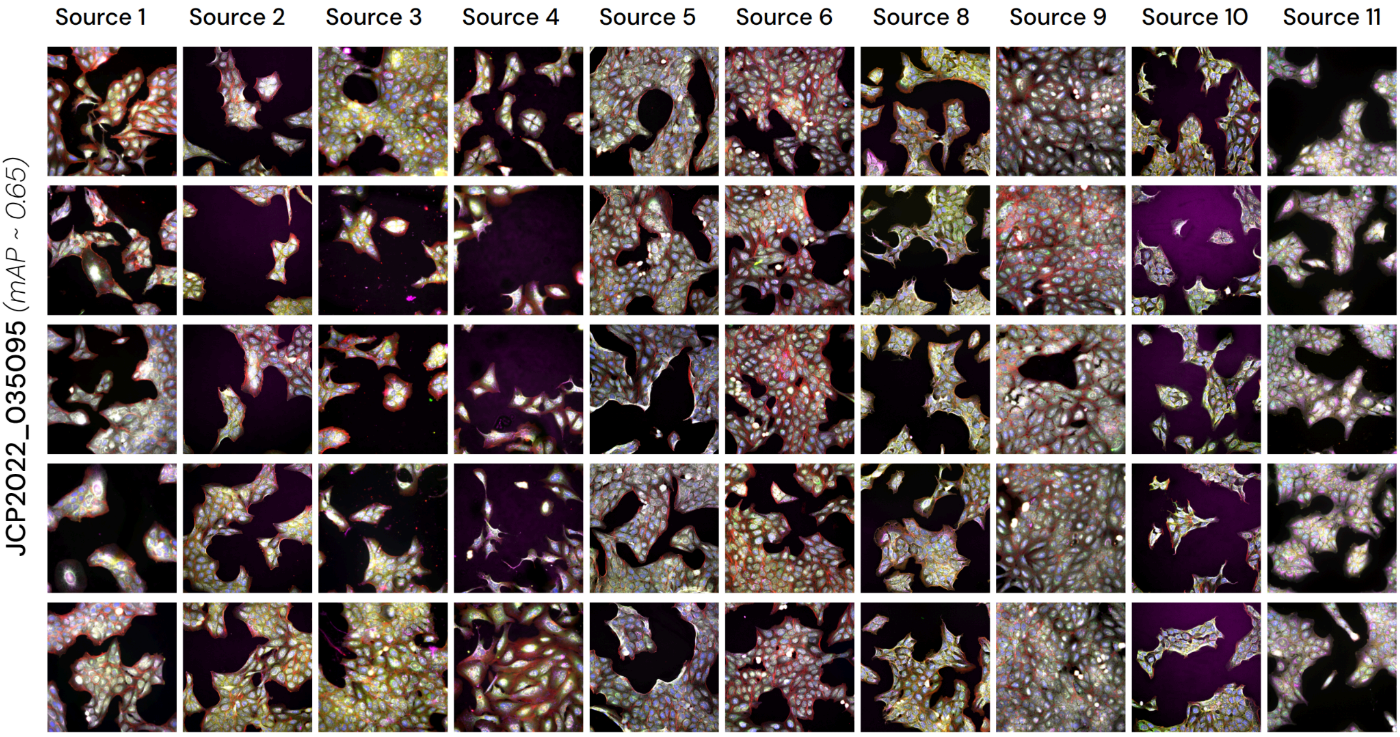
Source 1
Source 2
Source 3
Source 4
Source 5
Source 6
Source 8
Source 9
Source 10
Source 11
JCP2022_035095 (mAP ~ 0.65)

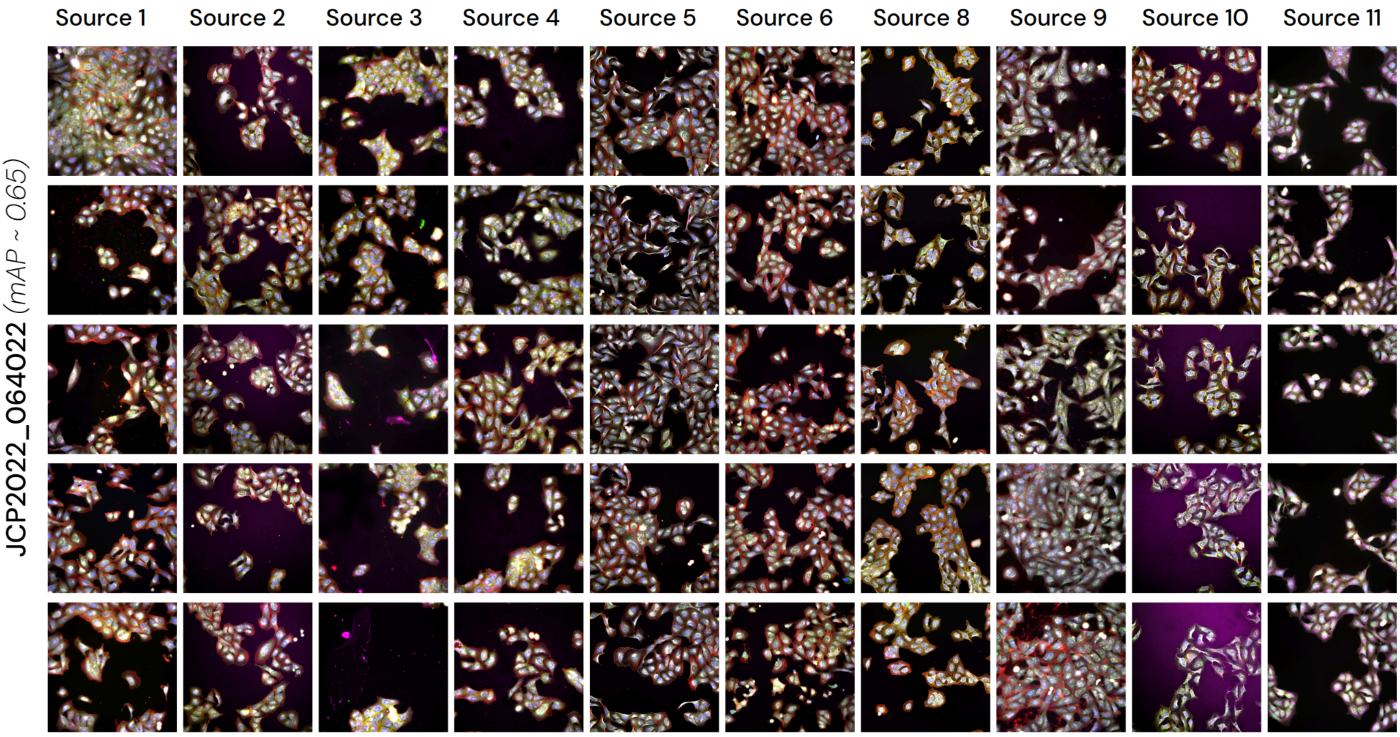
Source 1
Source 2
Source 3
Source 4
Source 5
Source 6
Source 8
Source 9
Source 10
Source 11
JCP2022_064022 (mAP ~ 0.65)

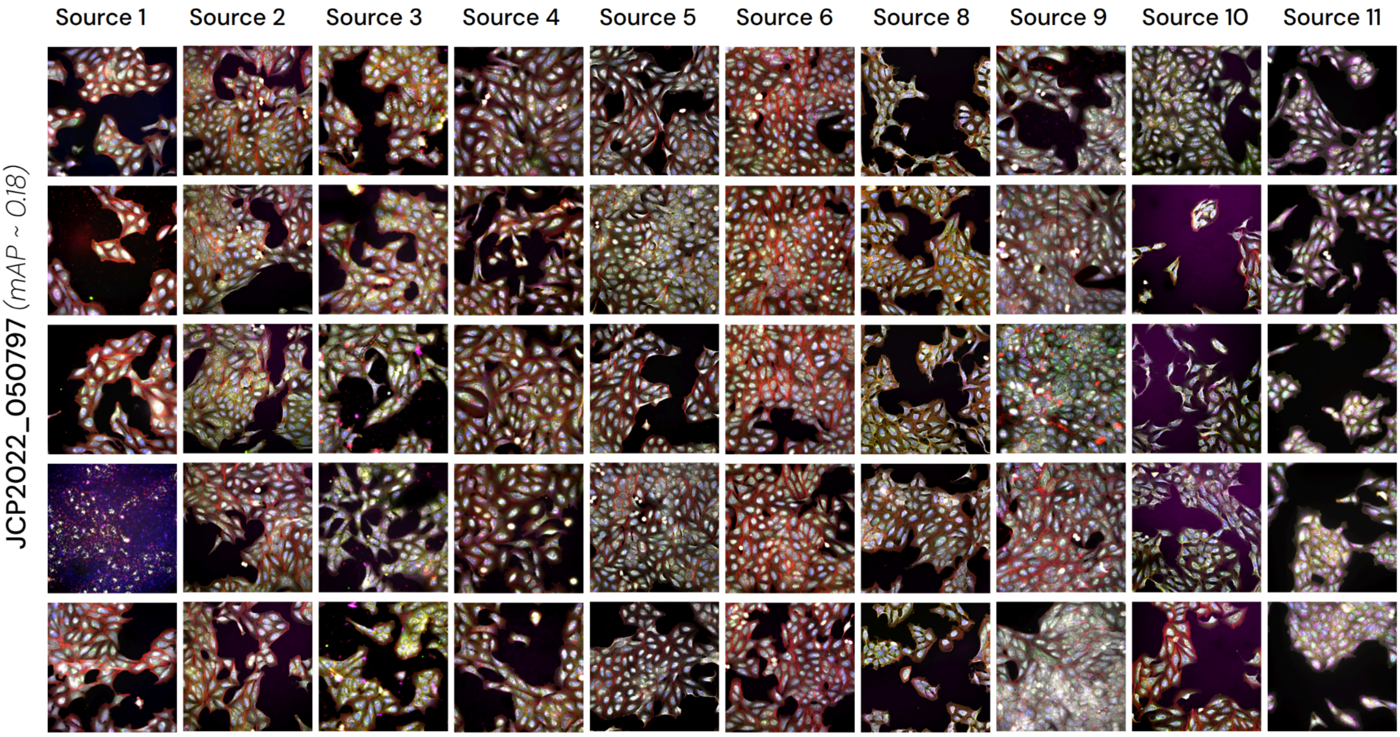
Source 1
Source 2
Source 3
Source 4
Source 5
Source 6
Source 8
Source 9
Source 10
Source 11
JCP2022_050797 (mAP ~ 0.18)

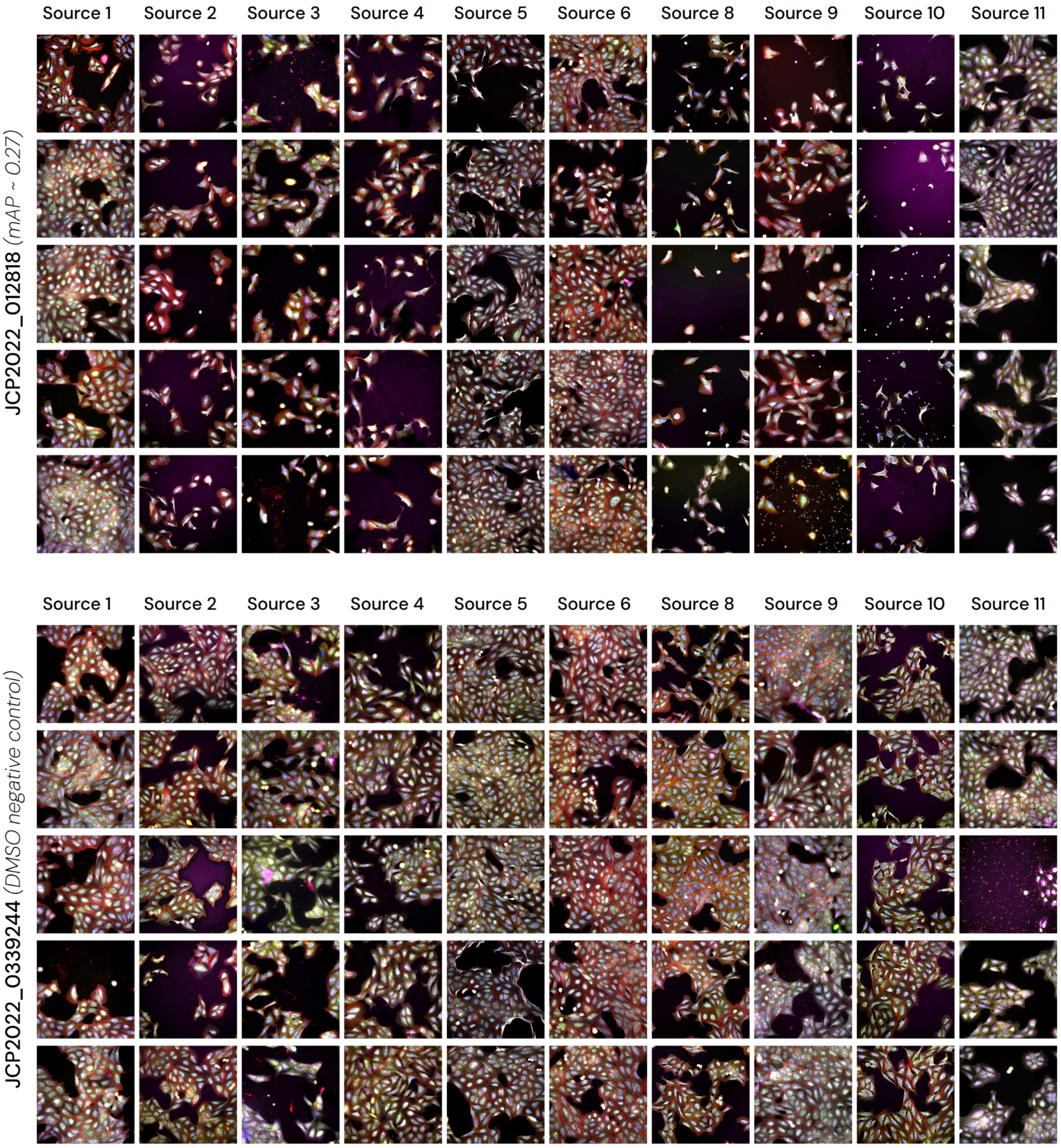


**Supplementary figure 2:** Random images from the JUMP-CP sources for each positive control compound. The last panel displays DMSO negative control samples for every source with its associated mAP value within our pipeline.

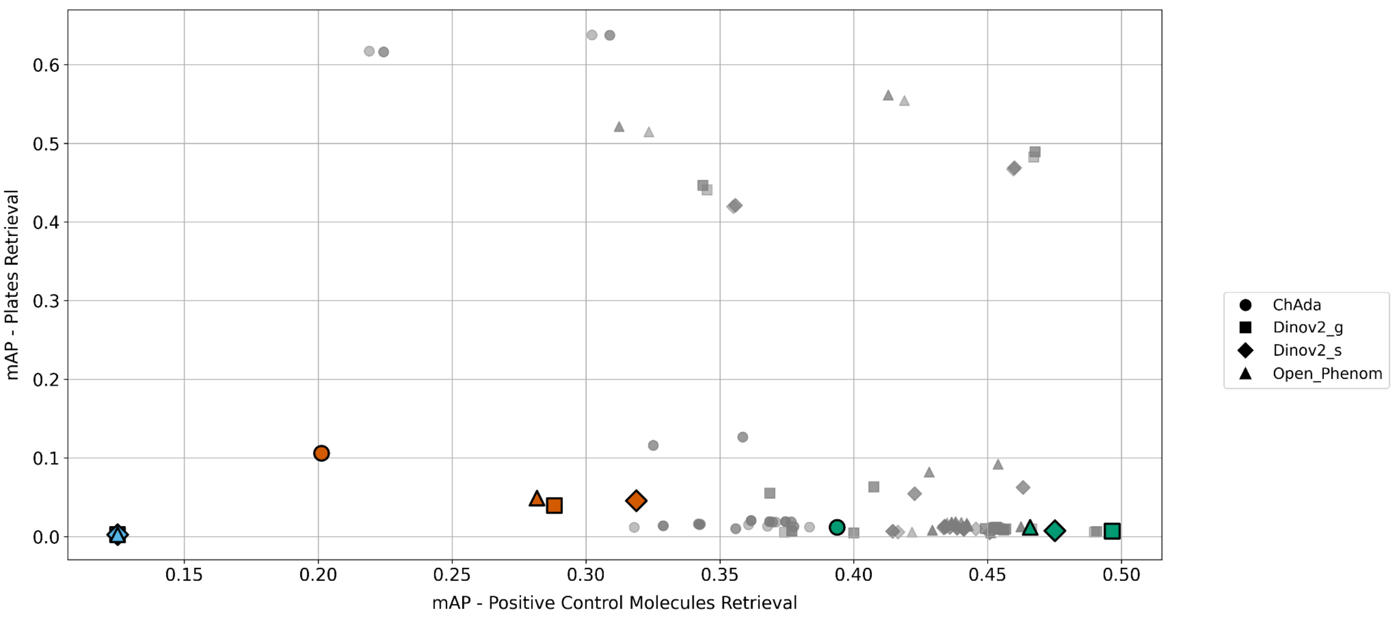


**Supplementary figure 3:** Sweep of 100 best normalization pipelines using control wells replicate samples from JUMP-CP. For each possible pipeline (represented on this plot by a single point), after processing all samples, we checked how good the pipeline was at retrieving sample replicates from the same perturbation (X axis, the higher the better) and at confounding replicate samples of negative control whatever the experimental plate (Y axis, the lower the better) by computing two mean Average Precisions (mAP) values

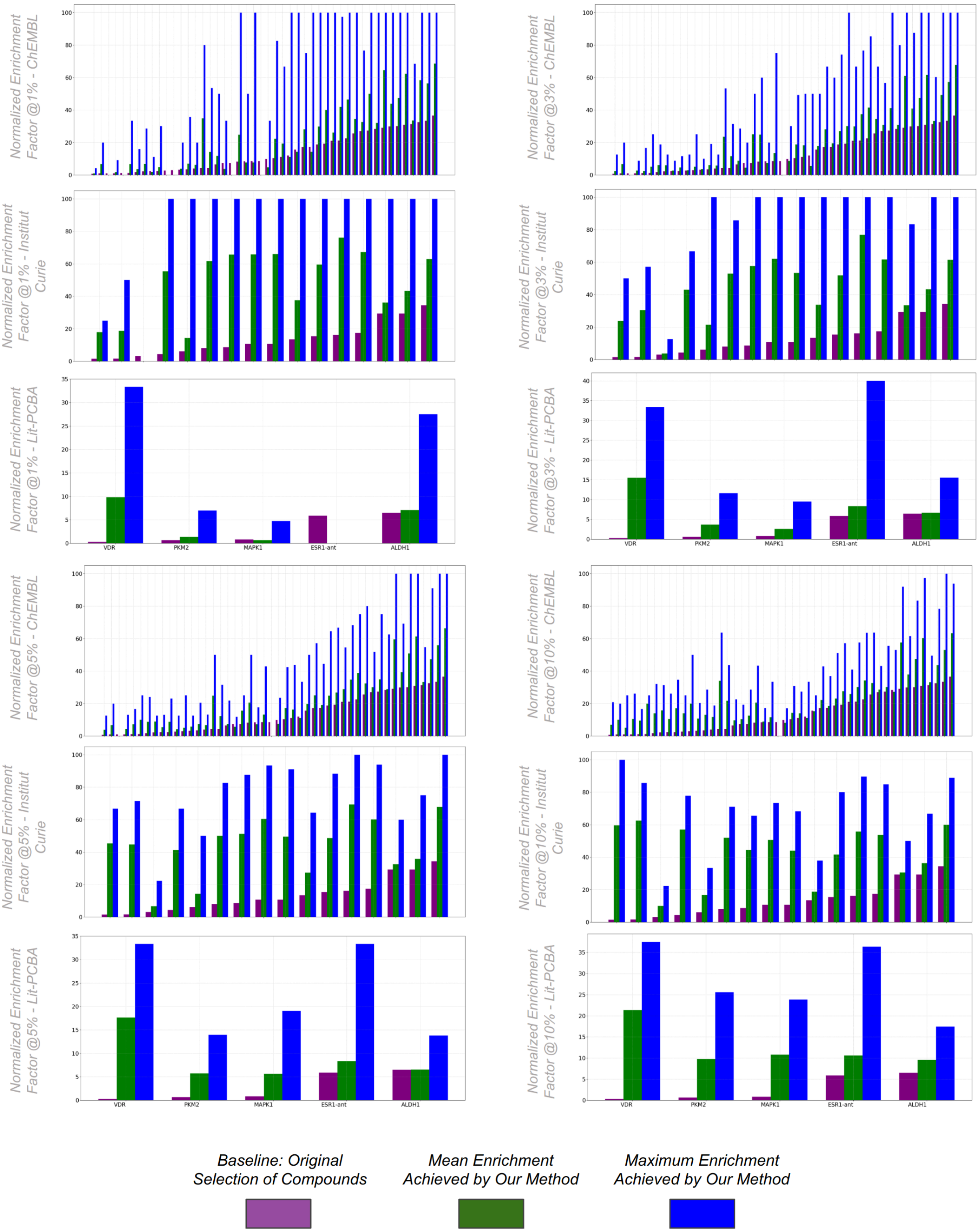


**Supplementary figure 4:** Same as **Figure 2** for a selection of 1, 3, 5 or 10% of all tested compounds for each screen.

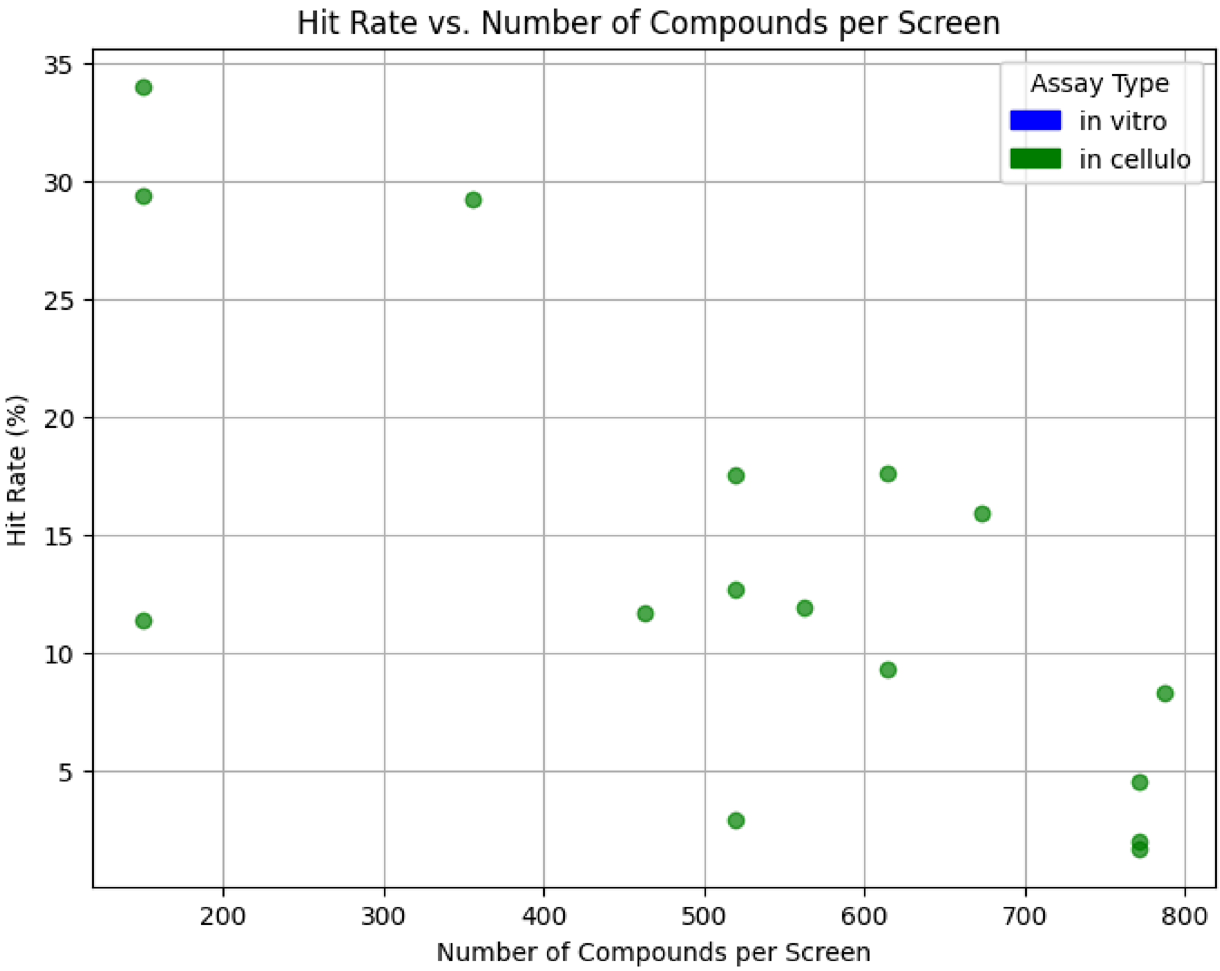


**Supplementary figure 5:** Compound counts and hit rates for the Institut Curie screens

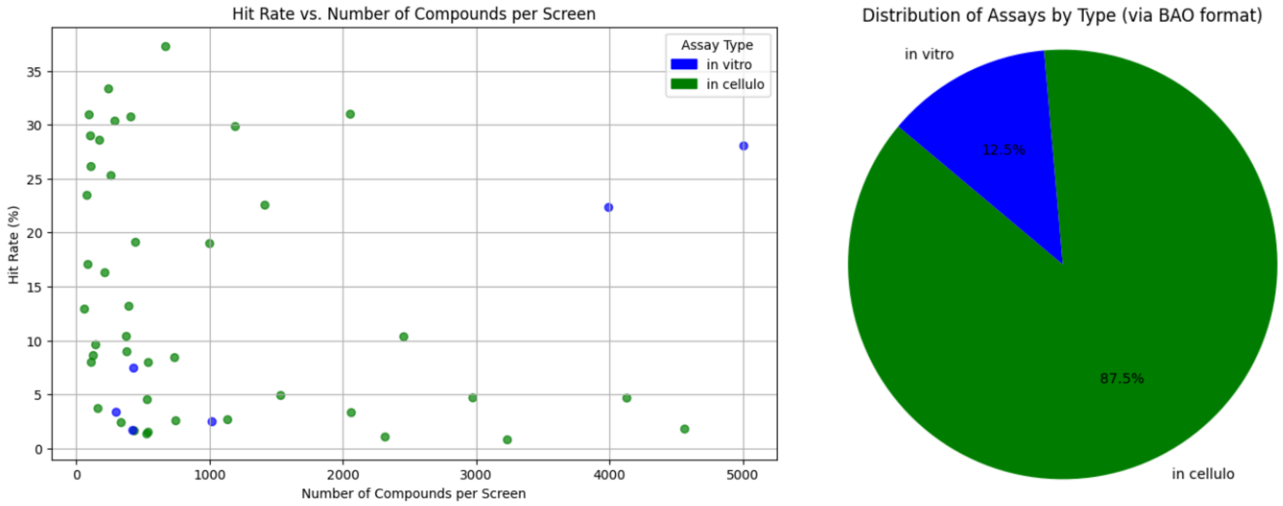


**Supplementary figure 6:** Compound counts, hit rates and assays types for ChEMBL screens

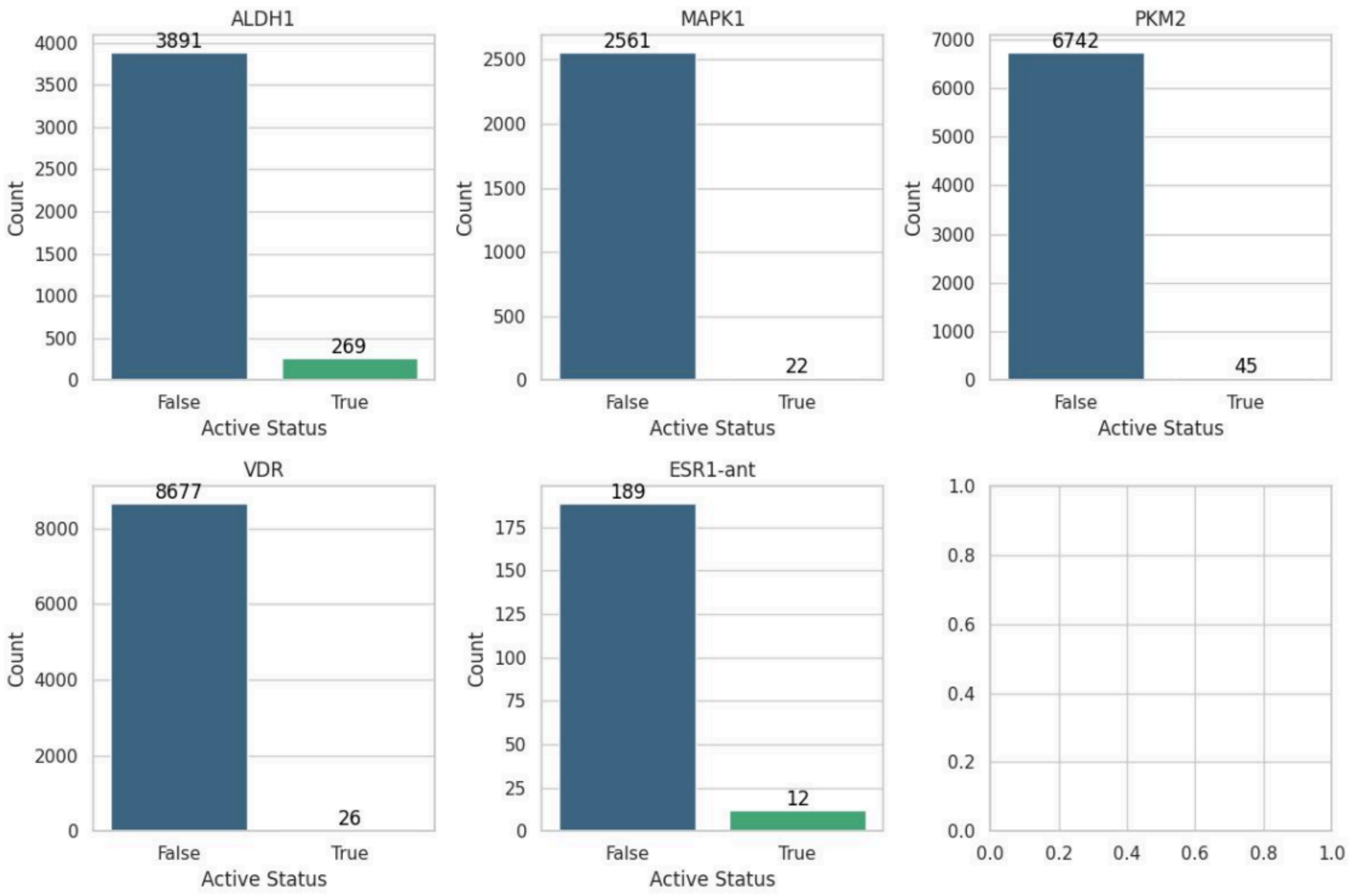


**Supplementary figure 7:** Distribution of active (green) and non active (blue) compounds that were both in the JUMP-CP and in the original Lit-PCBA benchmark for each target protein. Description and count for Lit-PCBA selected targets

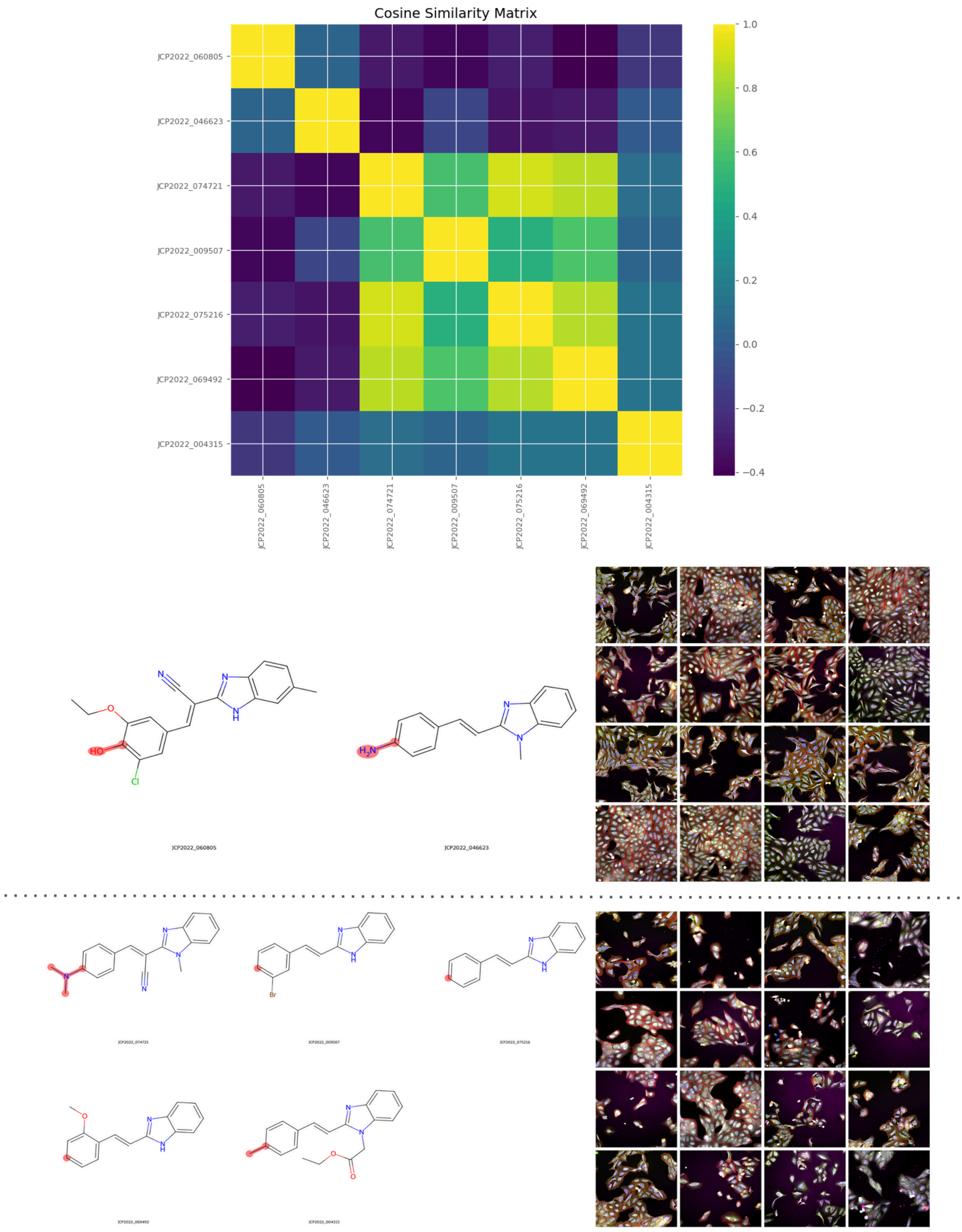


**Supplementary figure 8:** Example of a chemical series that displays phenotypic activity cliffs.

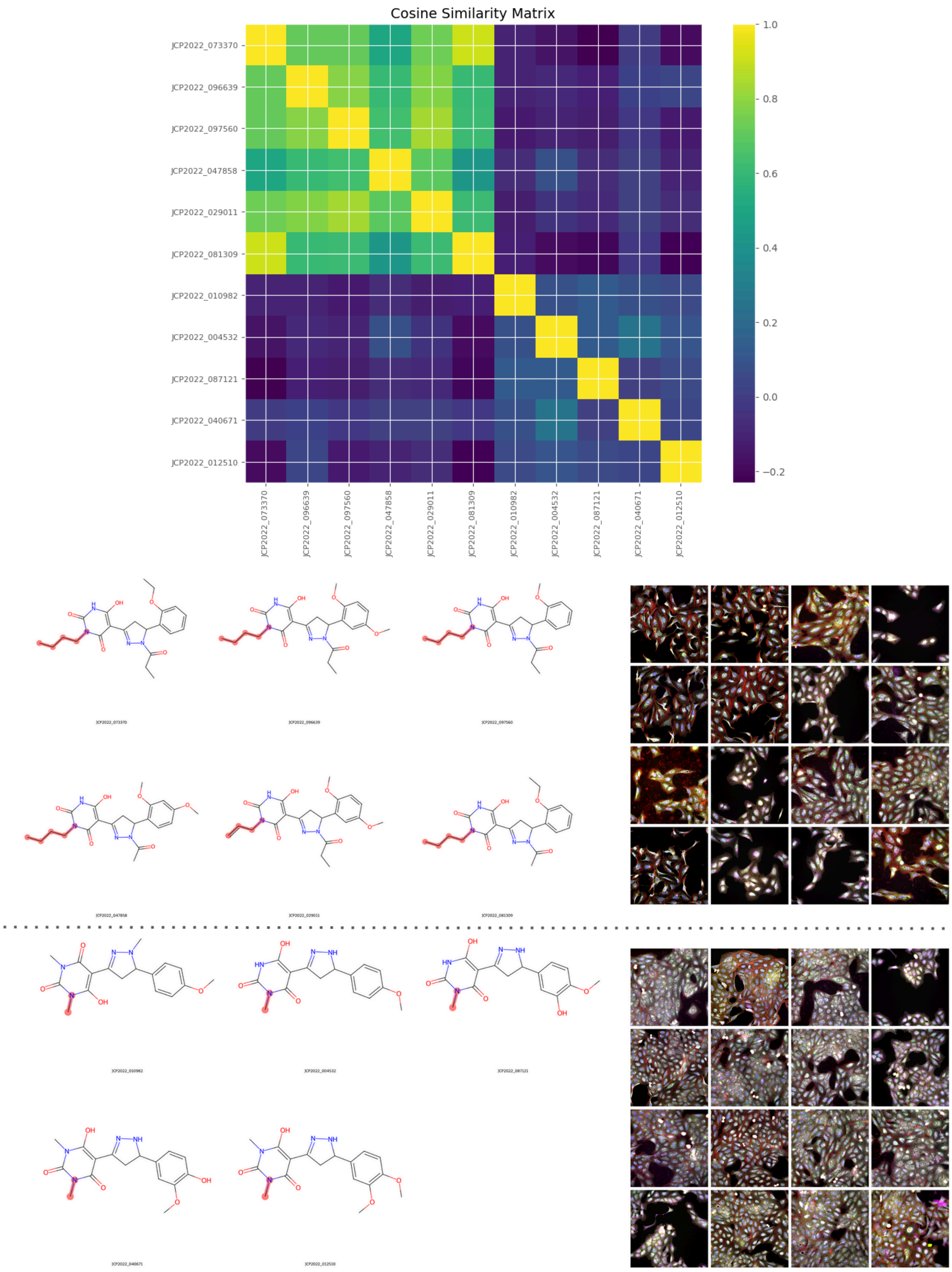


**Supplementary figure 9:** Example of a chemical series that displays phenotypic activity cliffs.

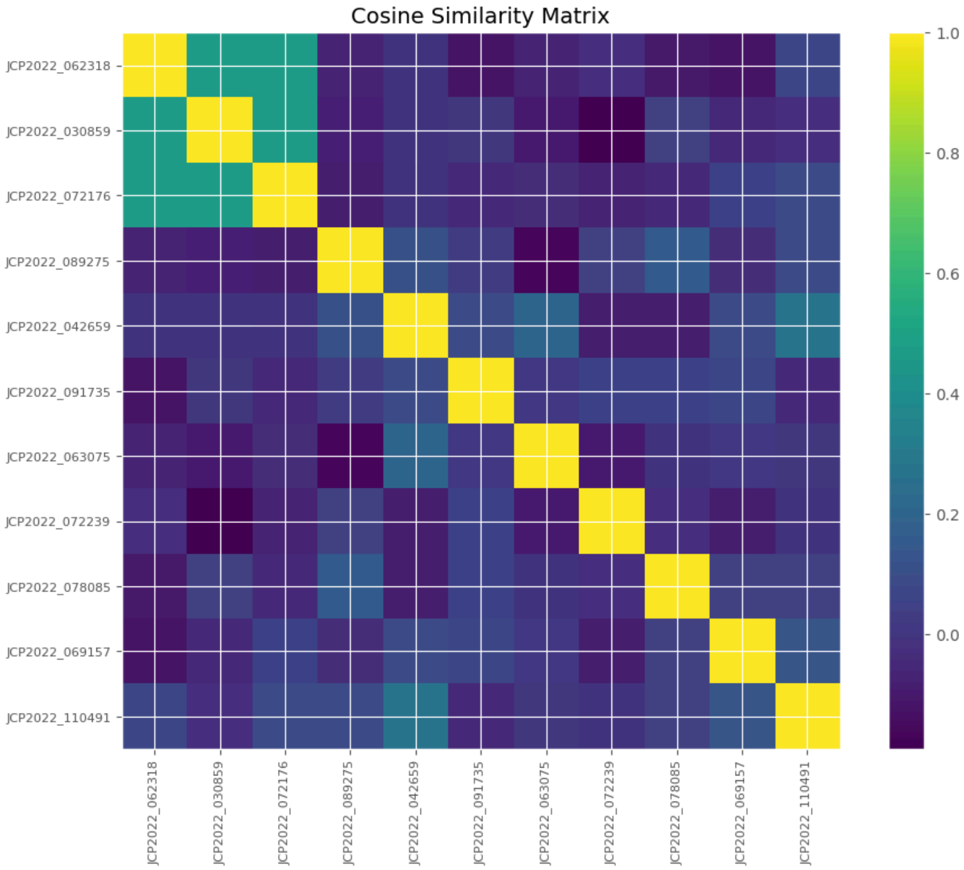
Cosine Similarity Matrix
JCP2022_062318
JCP2022_030859
JCP2022_072176
JCP2022_089275
JCP2022_042659
JCP2022_091735
JCP2022_063075
JCP2022_072239
JCP2022_078085
JCP2022_069157
JCP2022_110491
1.0
0.8
0.6
0.4
0.2
0.0

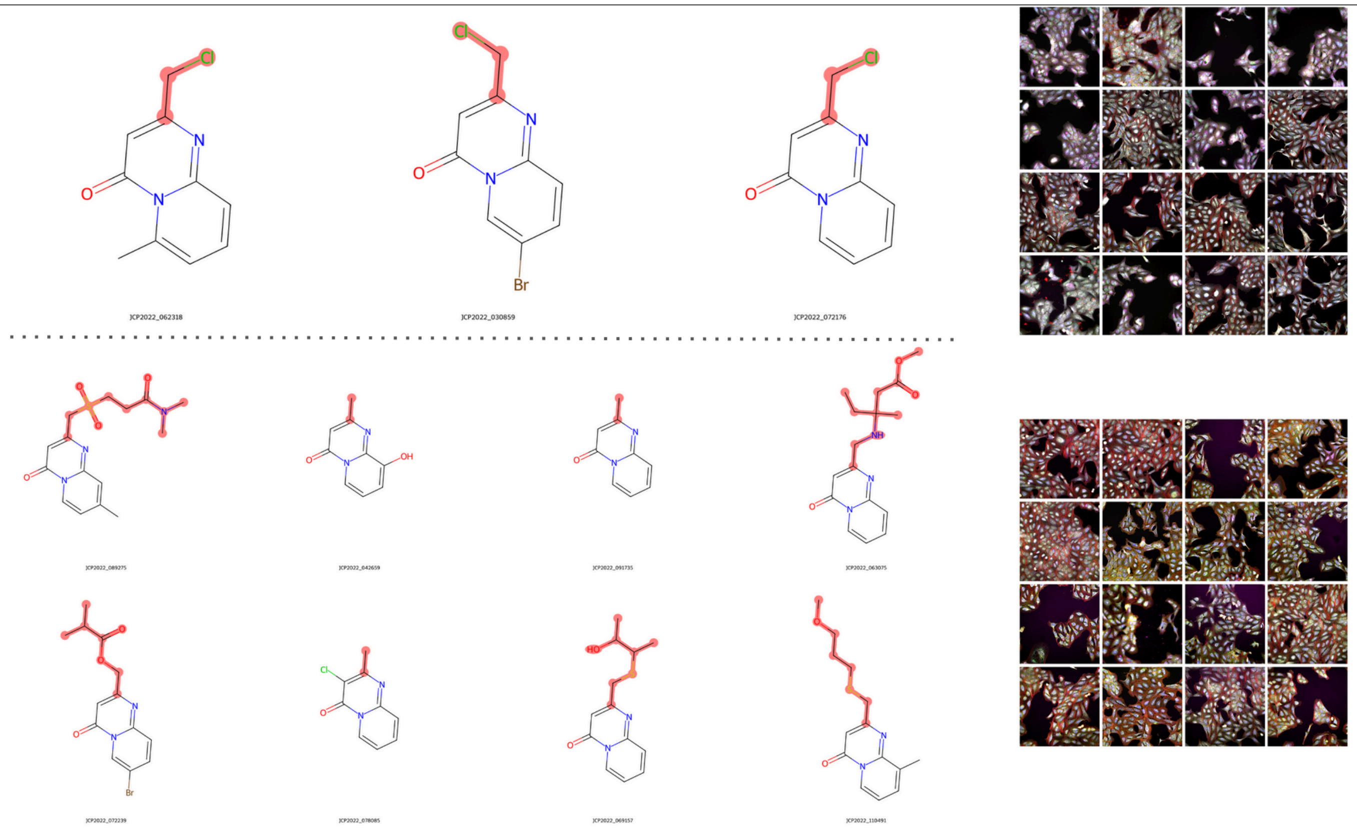
JCP2022_062318
JCP2022_030859
JCP2022_072176

**Supplementary figure 10:** Example of a chemical series that displays phenotypic activity cliffs.

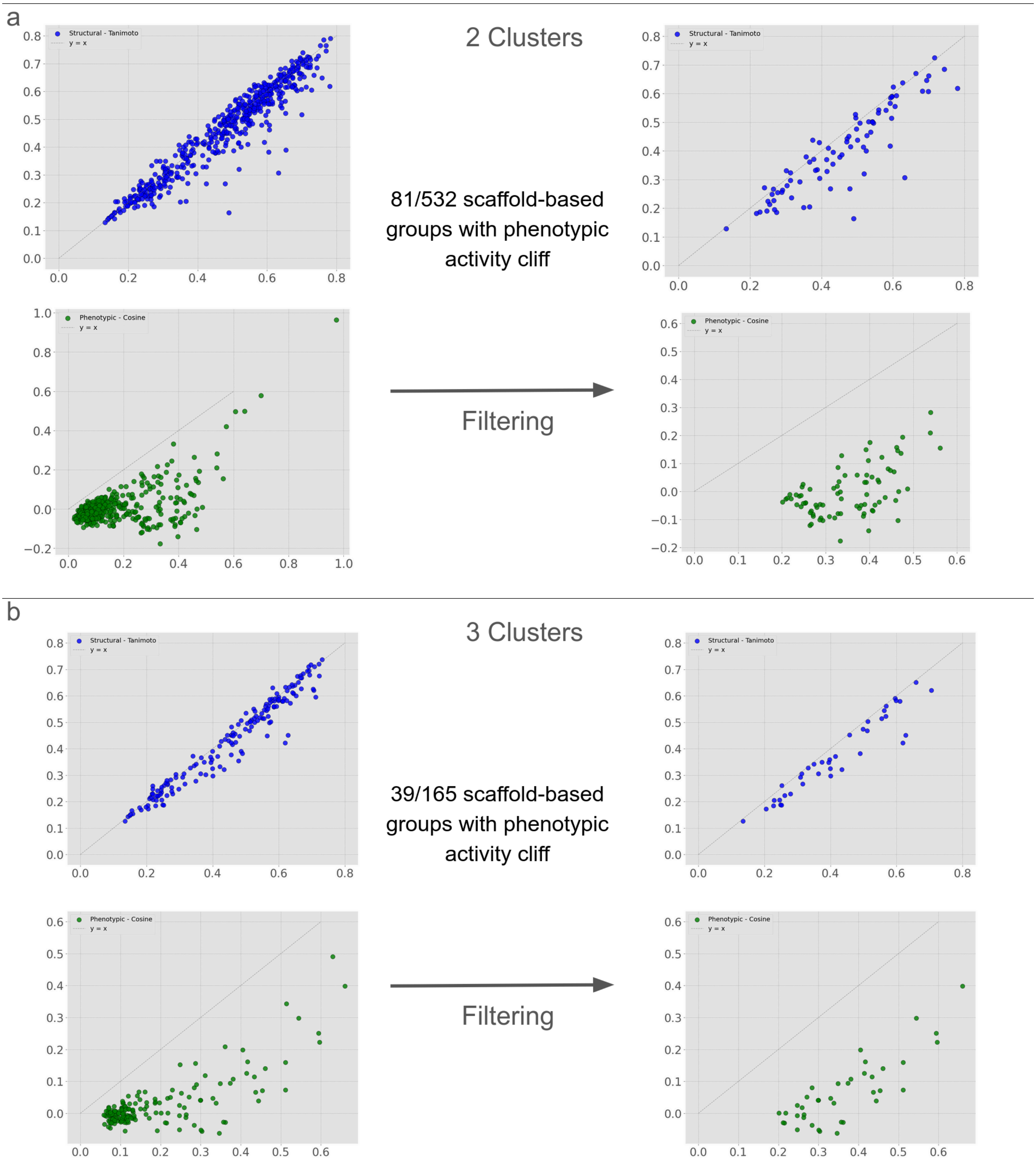

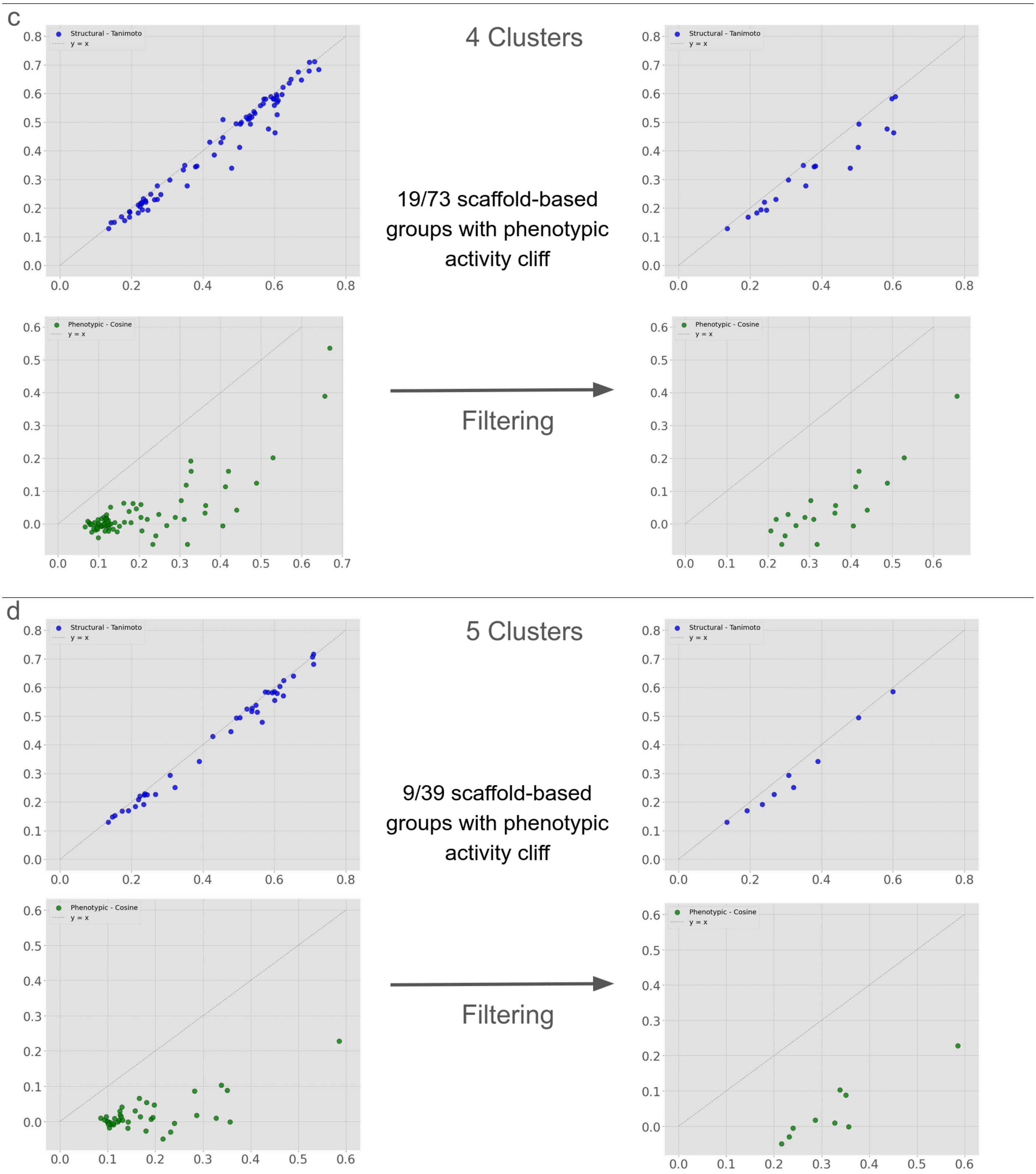


**Supplementary figure 11:** Same figure as **Figure 4** c and b for two, three, four, or five clusters per scaffold (2 being displayed in **Figure 4**)

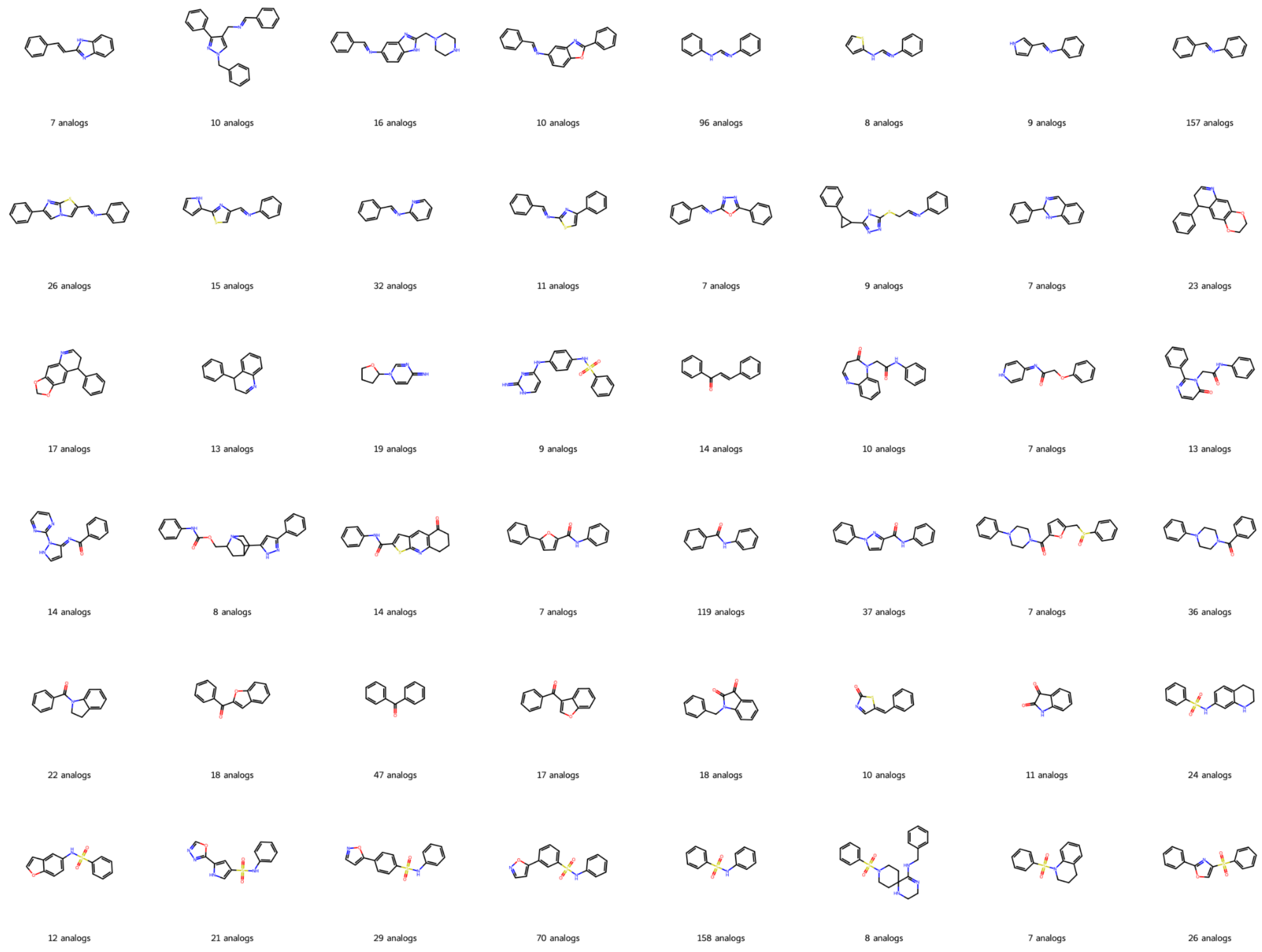


**Supplementary figure 12:** Example of Demis-Murcko scaffolds from series with phenotypic activity cliffs

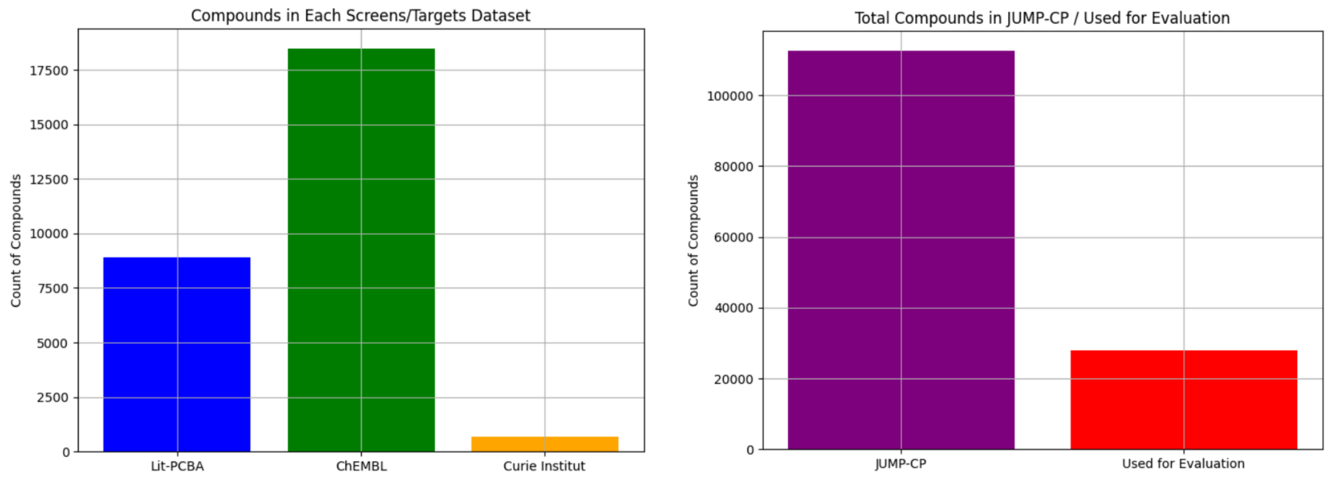


**Supplementary figure 13:** Proportion of the JUMP-CP dataset evaluated across all selected assays

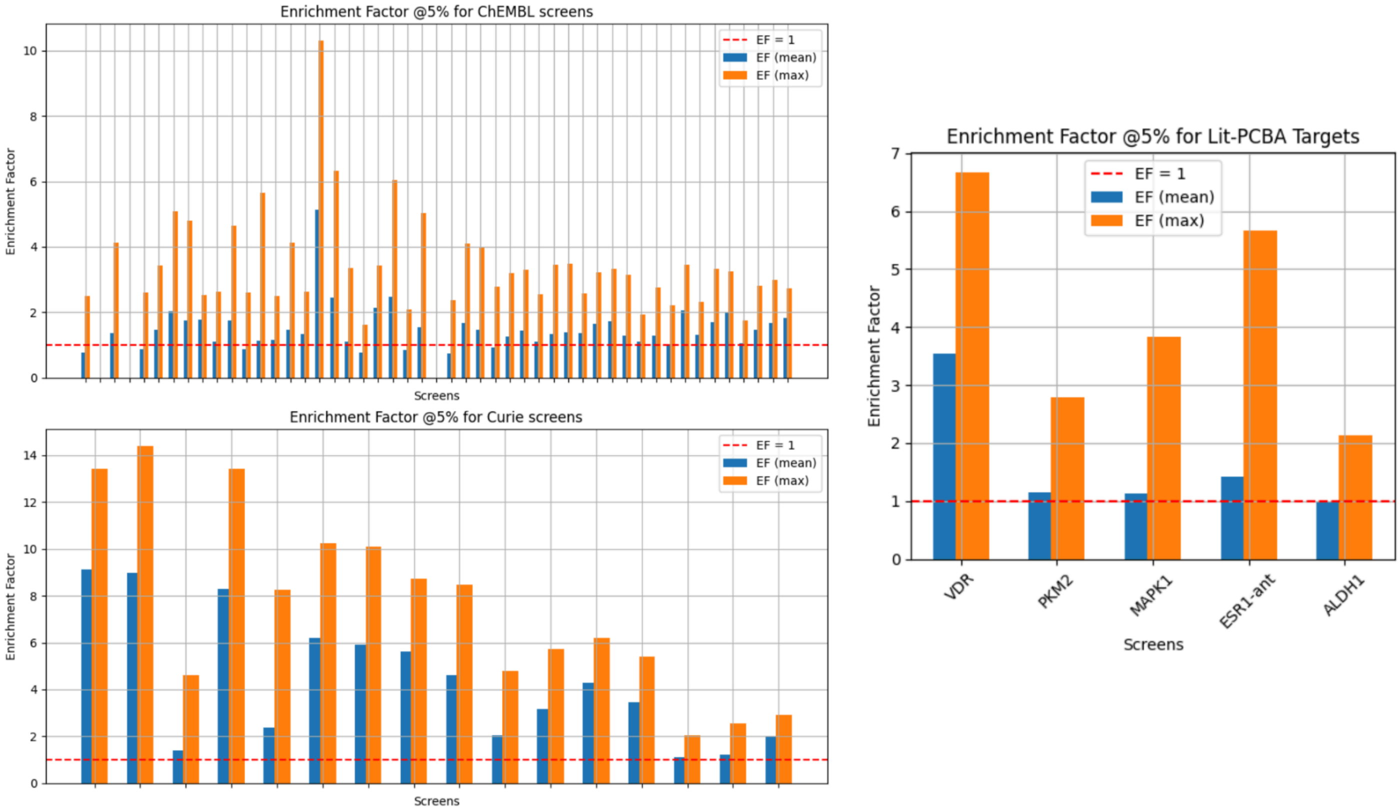


**Supplementary figure 14:** all enrichment factors at 5% for our method on every screen and target tested.

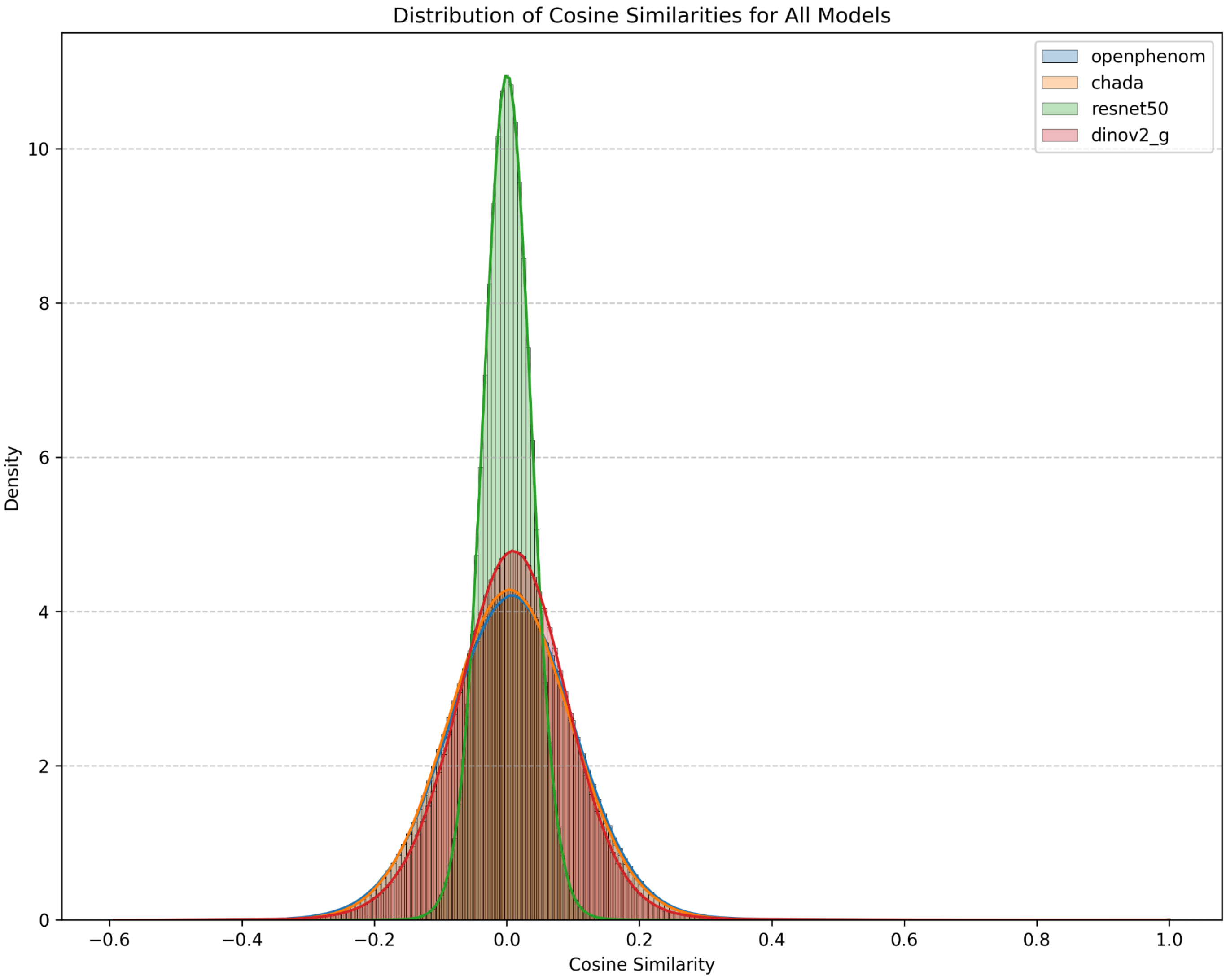


**Supplementary figure 15:** Distribution of all cosine similarities between all normalized phenotypic profiles of JUMP-CP molecules for each deep encoder.

### ChEMBL Screens

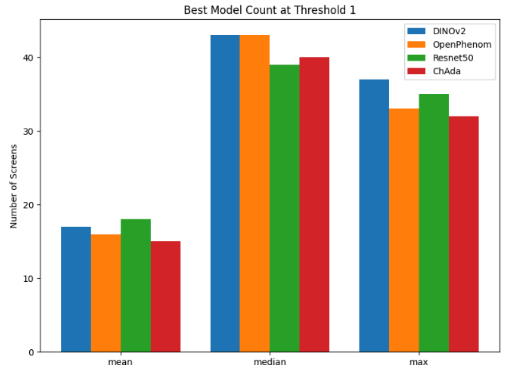


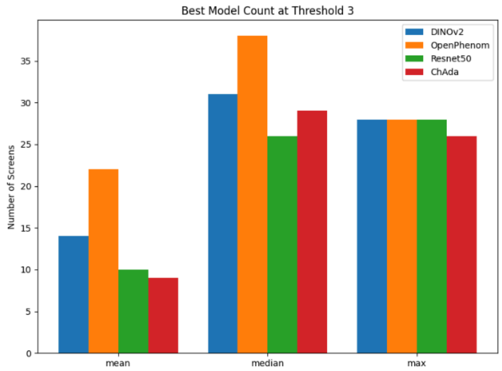


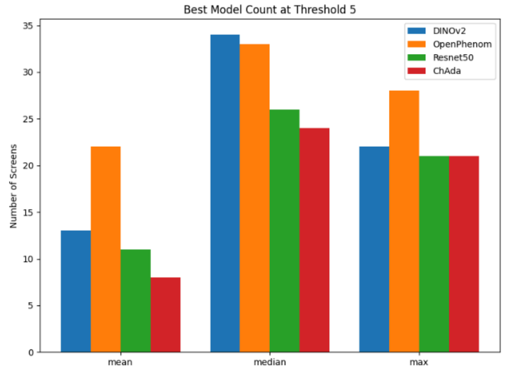


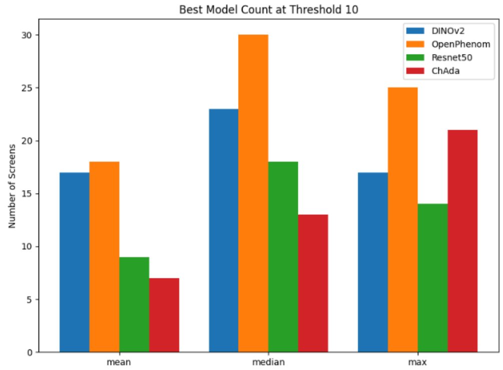


### Lit-PCBA

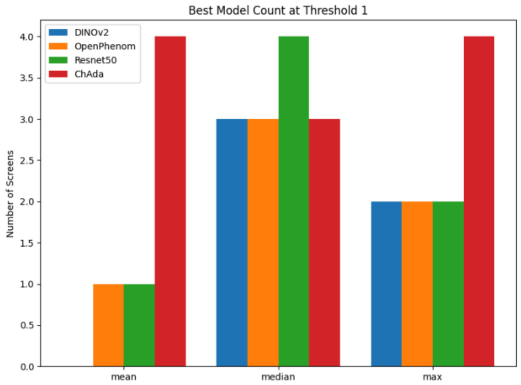


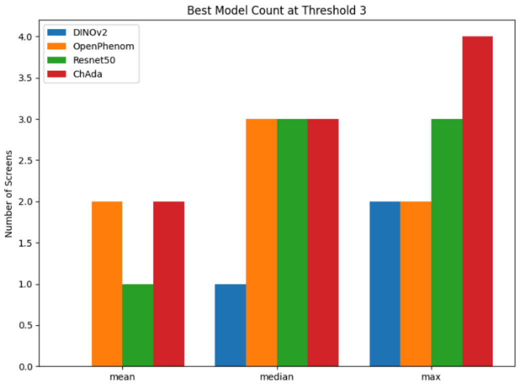


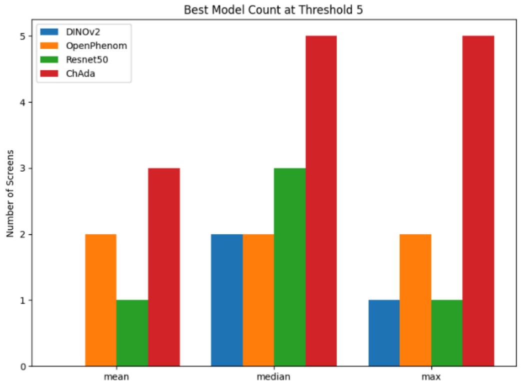


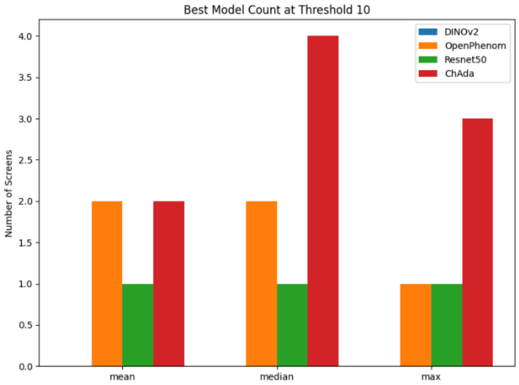

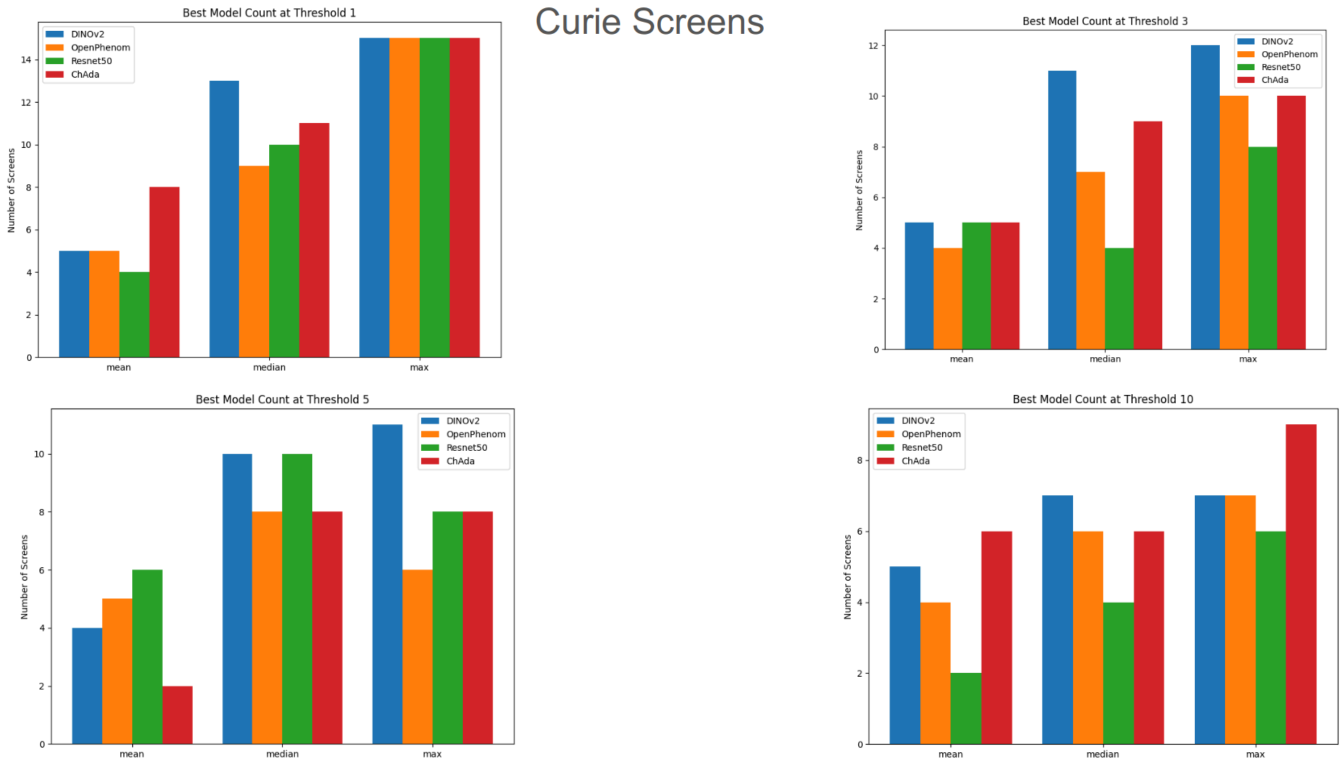


**Supplementary figure 16:** All results (EF, nEF, mean, median and max) for all percentage of selection are in csv files in https://github.com/mxfly14/phenoseeker/tree/main/results/. Here are displayed the number of screens and targets from every source where each encoder performs the best (Here DINOv2 is the Giant version). If two models perform the same, they are both counted.

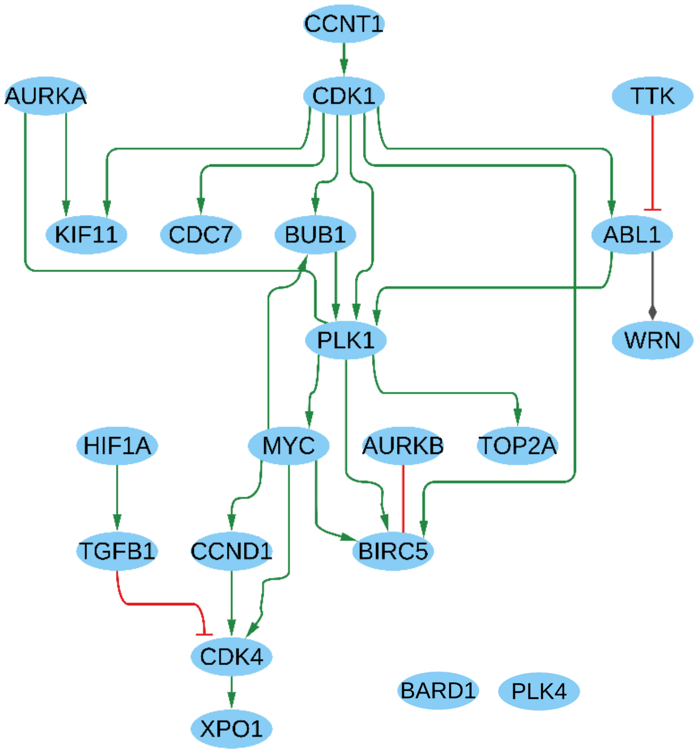


**Supplementary figure 17:** Network inferred from all the G2M genes targeted by a JUMP-CP compound. An extended version of this network could include intermediate genes.

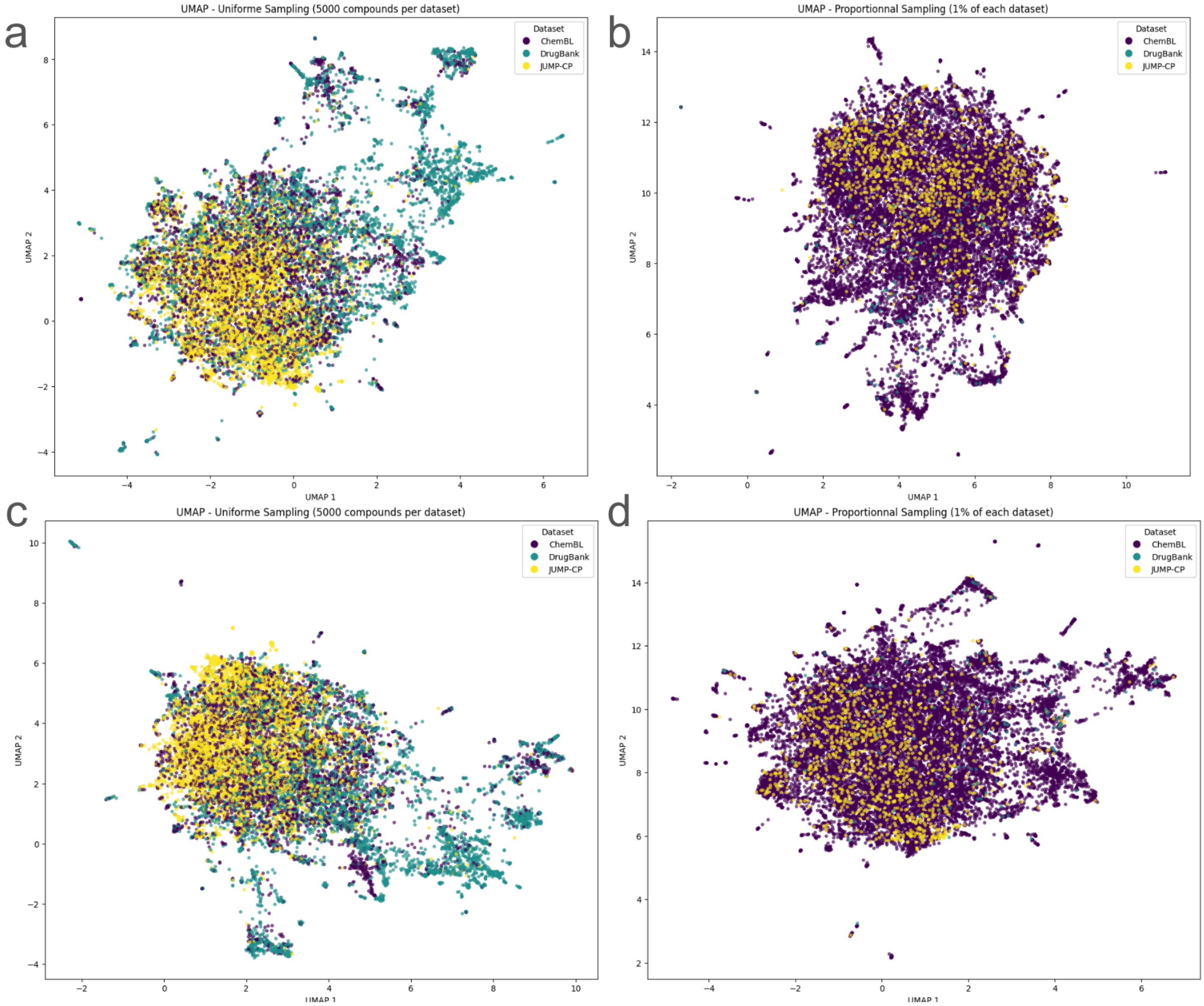


**Supplementary figure 18:** UMAP of Chemical space coverage for JUMP-CP versus drug bank and ChEMBL

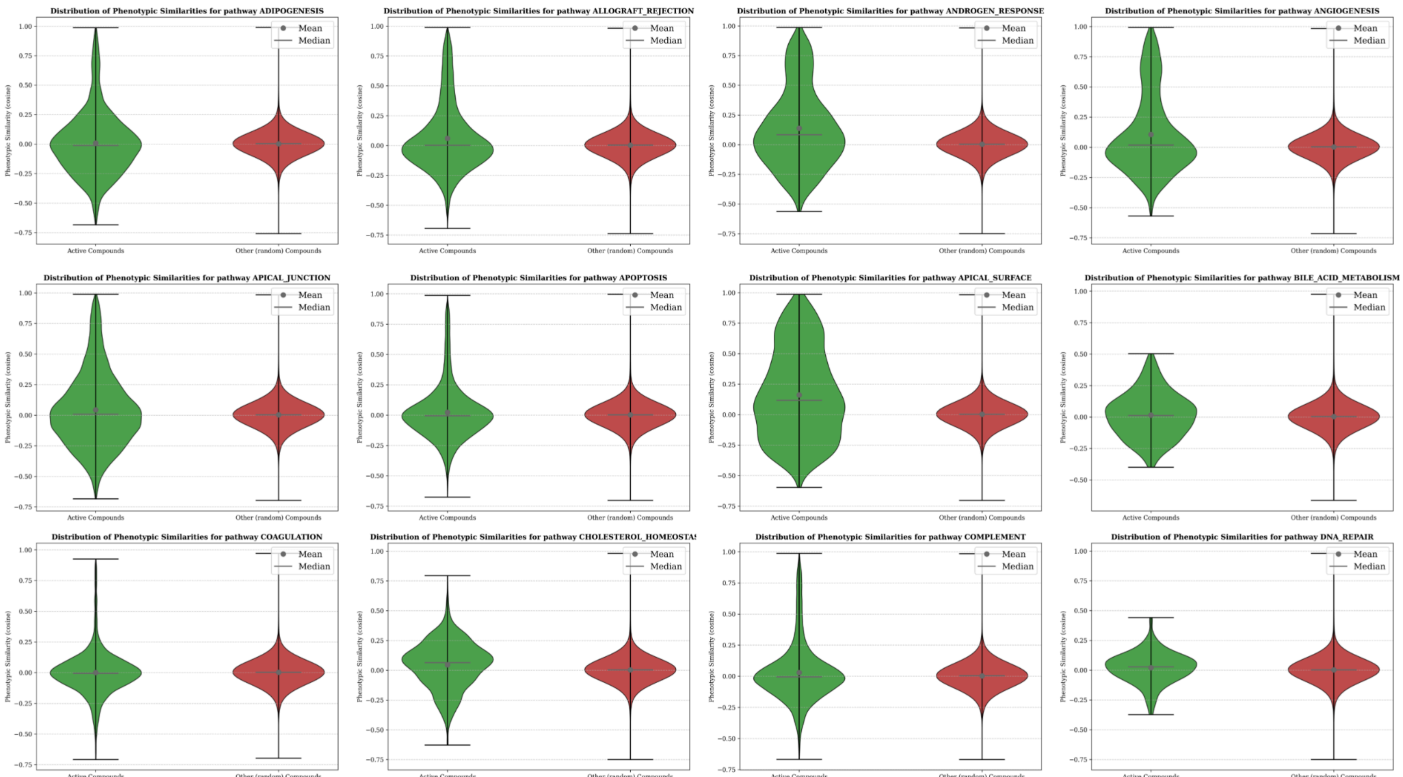
Distribution of Phenotypic Similarities for pathway ADIPOGENESIS
Distribution of Phenotypic Similarities for pathway ALLOGRAFT_REJECTION
Distribution of Phenotypic Similarities for pathway ANDROGEN_RESPONSE
Distribution of Phenotypic Similarities for pathway ANGIOGENESIS
Distribution of Phenotypic Similarities for pathway APICAL_JUNCTION
Distribution of Phenotypic Similarities for pathway APOPTOSIS
Distribution of Phenotypic Similarities for pathway APICAL_SURFACE
Distribution of Phenotypic Similarities for pathway BILE_ACID_METABOLISM
Distribution of Phenotypic Similarities for pathway COAGULATION
Distribution of Phenotypic Similarities for pathway CHOLESTEROL_HOMEOSTAS
Distribution of Phenotypic Similarities for pathway COMPLEMENT
Distribution of Phenotypic Similarities for pathway DNA_REPAIR
Mean
Median
Phenotypic Similarity (cosine)
Active Compounds
Other (random) Compounds

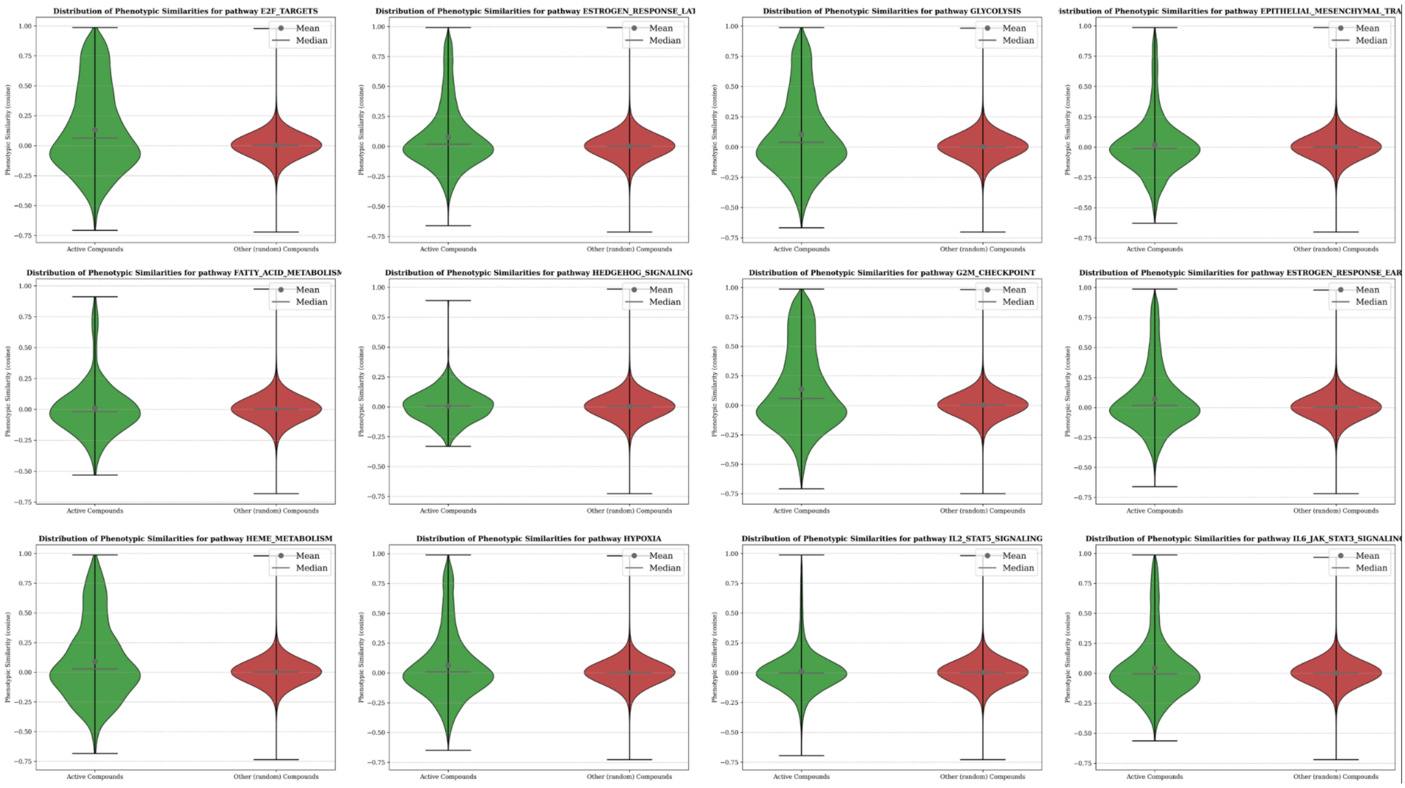
Distribution of Phenotypic Similarities for pathway E2F_TARGETS
Distribution of Phenotypic Similarities for pathway ESTROGEN_RESPONSE_LAT
Distribution of Phenotypic Similarities for pathway GLYCOLYSIS
istribution of Phenotypic Similarities for pathway EPITHELIAL_MESENCHYMAL_TRA
Distribution of Phenotypic Similarities for pathway FATTY_ACID_METABOLISM
Distribution of Phenotypic Similarities for pathway HEDGEHOG_SIGNALING
Distribution of Phenotypic Similarities for pathway G2M_CHECKPOINT
Distribution of Phenotypic Similarities for pathway ESTROGEN_RESPONSE_EAR
Distribution of Phenotypic Similarities for pathway HEME_METABOLISM
Distribution of Phenotypic Similarities for pathway HYPOXIA
Distribution of Phenotypic Similarities for pathway IL2_STAT5_SIGNALING
Distribution of Phenotypic Similarities for pathway IL6_JAK_STAT3_SIGNALING
Mean
Median
Phenotypic Similarity (cosine)
Active Compounds
Other (random) Compounds

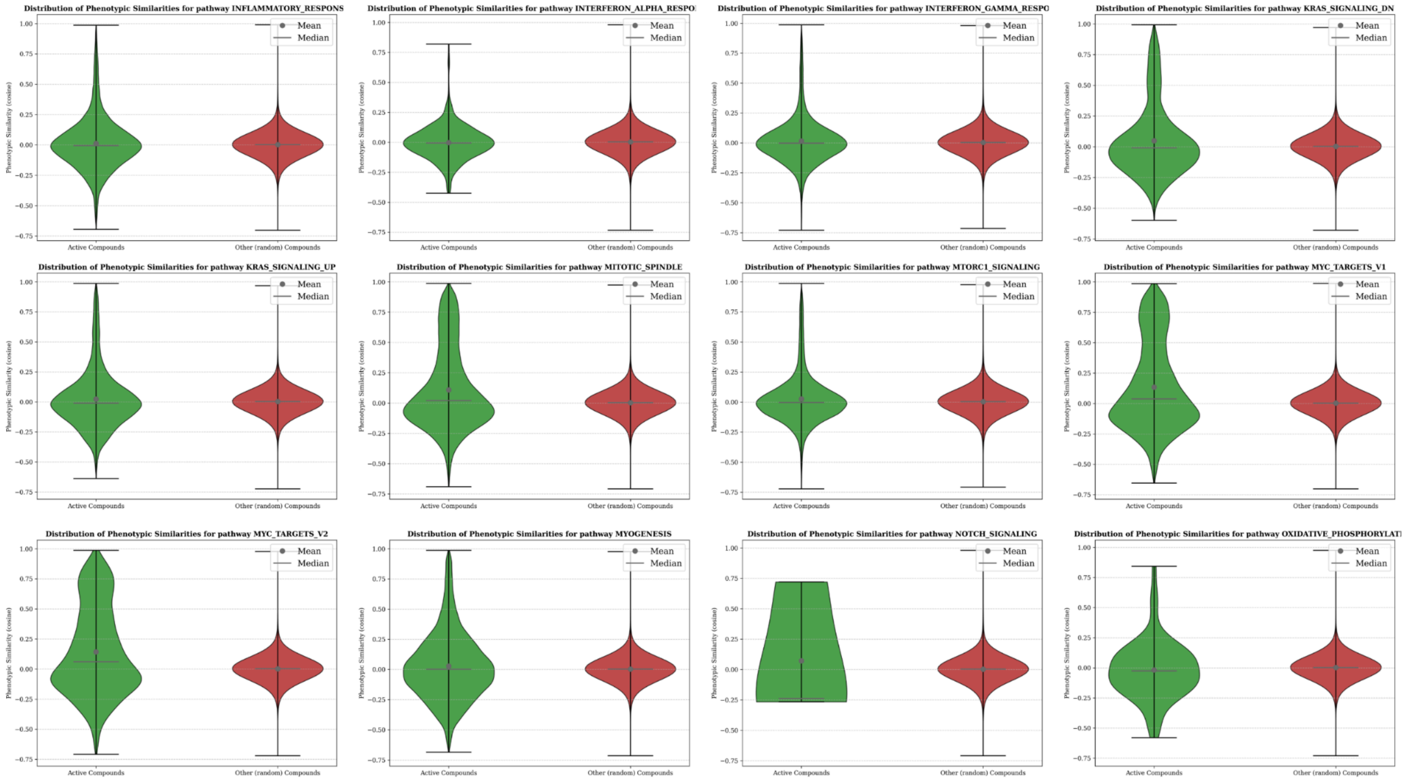
Distribution of Phenotypic Similarities for pathway INFLAMMATORY_RESPONS
Distribution of Phenotypic Similarities for pathway INTERFERON_ALPHA_RESPO
Distribution of Phenotypic Similarities for pathway INTERFERON_GAMMA_RESPO
Distribution of Phenotypic Similarities for pathway KRAS_SIGNALING_DN
Distribution of Phenotypic Similarities for pathway KRAS_SIGNALING_UP
Distribution of Phenotypic Similarities for pathway MITOTIC_SPINDLE
Distribution of Phenotypic Similarities for pathway MTORC1_SIGNALING
Distribution of Phenotypic Similarities for pathway MYC_TARGETS_V1
Distribution of Phenotypic Similarities for pathway MYC_TARGETS_V2
Distribution of Phenotypic Similarities for pathway MYOGENESIS
Distribution of Phenotypic Similarities for pathway NOTCH_SIGNALING
Distribution of Phenotypic Similarities for pathway OXIDATIVE_PHOSPHORYLATI
Phenotypic Similarity (cosine)
Active Compounds
Other (random) Compounds
Mean
Median

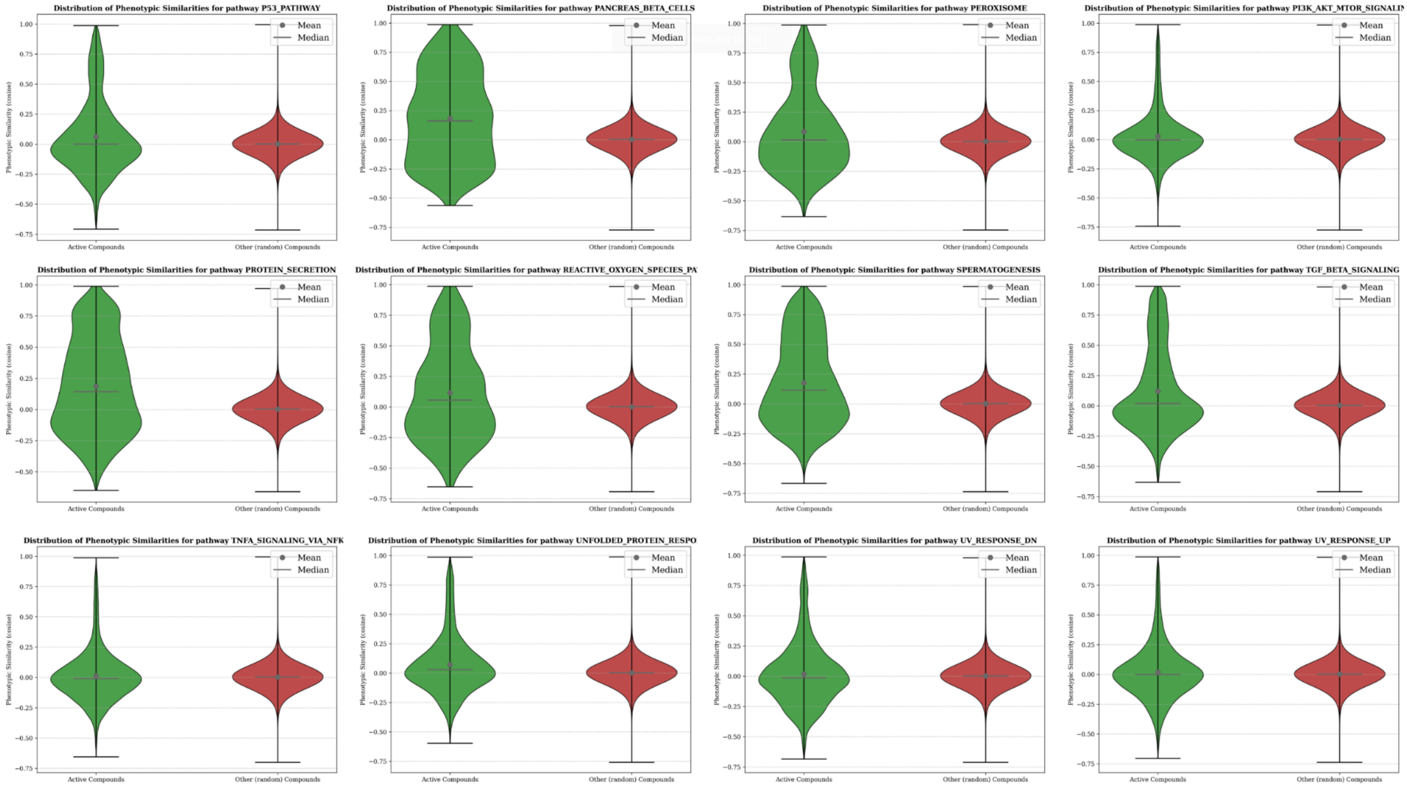
Distribution of Phenotypic Similarities for pathway P53_PATHWAY
Distribution of Phenotypic Similarities for pathway PANCREAS_BETA_CELLS
Distribution of Phenotypic Similarities for pathway PEROXISOME
Distribution of Phenotypic Similarities for pathway PI3K_AKT_MTOR_SIGNALIN
Distribution of Phenotypic Similarities for pathway PROTEIN_SECRETION
Distribution of Phenotypic Similarities for pathway REACTIVE_OXYGEN_SPECIES_PA
Distribution of Phenotypic Similarities for pathway SPERMATOGENESIS
Distribution of Phenotypic Similarities for pathway TGF_BETA_SIGNALING
Distribution of Phenotypic Similarities for pathway TNFA_SIGNALING_VIA_NFK
Distribution of Phenotypic Similarities for pathway UNFOLDED_PROTEIN_RESPO
Distribution of Phenotypic Similarities for pathway UV_RESPONSE_DN
Distribution of Phenotypic Similarities for pathway UV_RESPONSE_UP
Phenotypic Similarity (cosine)
Active Compounds
Other (random) Compounds
Mean
Median

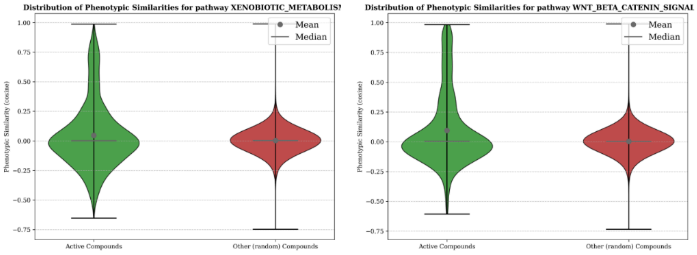


**Supplementary figure 19:** Distribution of phenotypic similarities (based on cosine similarities) for every pathway of the Hallmarks of Cancer gene sets

### Supplementary Tables

| Perturbation Name | Metadata_JCP2022 ID | InChIKey | Vendor URL |
|---|---|---|---|
| AMG900 | JCP2022_037716 | IVUGFMLRJOCGAS-UHFFFAOYSA-N | https://www.selleckchem.com/products/amg-900.html |
| NVS-PAK1-1 | JCP2022_064022 | OINGHOPGNMYCAB-INIZCTEOSA-N | https://www.medchemexpress.com/NVS-PAK1-1.html |
| dexamethasone | JCP2022_025848 | UREBDLICKHMUKA-CXSFZGCWSA-N | https://www.tocris.com/products/dexamethasone_1126 |
| LY2109761 | JCP2022_035095 | IHLVSLOZUHKNMQ-UHFFFAOYSA-N | https://www.selleckchem.com/products/ly2109761.html |
| FK-866 | JCP2022_046054 | KPBNHDGDUADAGP-VAWYXSNFSA-N | https://www.selleckchem.com/products/apo866-fk866.html |
| quinidine | JCP2022_050797 | LOUPRKONTZGTKE-LHHVKLHASA-N | https://www.medchemexpress.com/Quinidine.html |
| TC-S7004 | JCP2022_012818 | CQKBSRPVZZLCJE-UHFFFAOYSA-N | https://www.tocris.com/products/tc-s-7004_5088 |
| Aloxistatin | JCP2022_085227 | SRVFFFJZQVENJC-IHRRRGAJSA-N | https://www.medchemexpress.com/Aloxistatin.html |

**Supplementary Table 1:** List of positive control compounds. More details on selection of those compounds and their experimental plate distribution can be found here : [https://github.com/jump-cellpainting/JUMP-Target?tab=readme-ov-file](https://github.com/jump-cellpainting/JUMP-Target?tab=readme-ov-file)

| Source | Comparison | Normality p-value | Test Used | Test Statistic | p-value | Cohen's d |
|---|---|---|---|---|---|---|
| Institut Curie | Top 5% Structure vs. All | 0.8522102299 | Paired t-test | 21.1972672683 | 0.0000000000 | 5.1209 |
| | Top 5% Structure vs. Top 5% Phenotypic | 0.0084530672 | Wilcoxon signed-rank test | 136.0000000000 | 0.0002184045 | 3.5990 |
| | Top 5% Phenotypic vs. All | 0.4981551354 | Paired t-test | 1.2699852770 | 0.1117189245 | 0.3587 |
| Chembl | Top 5% Structure vs. All | 0.2436110830 | Paired t-test | 23.2947313786 | 0.0000000000 | 3.1605 |
| | Top 5% Structure vs. Top 5% Phenotypic | 0.0125169596 | Wilcoxon signed-rank test | 1225.0000000000 | 0.0000000000 | 2.9893 |
| | Top 5% Phenotypic vs. All | 0.0000286625 | Wilcoxon signed-rank test | 549.0000000000 | 0.7361682603 | 0.0319 |
| JUMP-CP | Top 5% Structure vs. All | 0.0000000194 | Wilcoxon signed-rank test | 50005000.0000 | 0.0000000000 | 6.1630 |
| | Top 5% Structure vs. Top 5% Phenotypic | 0.0000000281 | Wilcoxon signed-rank test | 50005000.0000 | 0.0000000000 | 5.5920 |
| | Top 5% Phenotypic vs. All | 0.0000000000 | Wilcoxon signed-rank test | 49883931.0000 | 0.0000000000 | 0.9341 |

**Supplementary Table 2:** Statistical details for **Figure 3**

| Role in Fig. 3 | Metadata_JCP2022 | Metadata_InChI |
|---|---|---|
| Positive Control (Red) | JCP2022_021041 | InChI=1S/C24H26FNO4/c1-15(2)26-21-6-4-3-5-20(21)24(16-7-9-17(25)10-8-16)22(26)12-11-18(27)13-19(28)14-23(29)30/h3-12,15,18-19,27-28H,13-14H2,1-2H3,(H,29,30) |
| Selected Compound (Blue left) | JCP2022_077096 | InChI=1S/C12H15N3O3/c1-3-6-18-8-4-5-9-10(7-8)14-11(13-9)15-12(16)17-2/h4-5,7H,3,6H2,1-2H3,(H2,13,14,15,16) |
| Selected Compound (Blue center) | JCP2022_007041 | InChI=1S/C59H84N18O14/c1-31(2)22-40(49(82)68-39(12-8-20-64-57(60)61)56(89)77-21-9-13-46(77)55(88)75-76-58(62)90)69-54(87)45(29-91-59(3,4)5)74-50(83)41(23-32-14-16-35(79)17-15-32)70-53(86)44(28-78)73-51(84)42(24-33-26-65-37-11-7-6-10-36(33)37)71-52(85)43(25-34-27-63-30-66-34)72-48(81)38-18-19-47(80)67-38/h6-7,10-11,14-17,26-27,30-31,38-46,65,78-79H,8-9,12-13,18-25,28-29H2,1-5H3,(H,63,66)(H,67,80)(H,68,82)(H,69,87)(H,70,86)(H,71,85)(H,72,81)(H,73,84)(H,74,83)(H,75,88)(H4,60,61,64)(H3,62,76,90) |
| Selected Compound (Blue right) | JCP2022_057581 | InChI=1S/C18H22N2O/c1-17(2,3)14-8-12(7-13(10-19)11-20)9-15(16(14)21)18(4,5)6/h7-9,21H,1-6H3 |
| Selected Compound (Turquoise left) | JCP2022_112877 | InChI=1S/C18H29N3O6/c1-4-8-19-15(22)13-14(27-13)16(23)20-12(10(3)5-2)17(24)21-9-6-7-11(21)18(25)26/h10-14H,4-9H2,1-3H3,(H,19,22)(H,20,23)(H,25,26) |
| Not Selected (Turquoise right) | JCP2022_103583 | InChI=1S/C19H31N3O6/c1-5-9-20-16(23)14-15(28-14)17(24)21-13(11(3)6-2)18(25)22-10-7-8-12(22)19(26)27-4/h11-15H,5-10H2,1-4H3,(H,20,23)(H,21,24) |
| Selected Compound (Orange left) | JCP2022_013346 | InChI=1S/C15H12N2O2/c16-15(19)17-12-7-3-1-5-10(12)9-14(18)11-6-2-4-8-13(11)17/h1-8H,9H2,(H2,16,19) |
| Not Selected (Orange right) | JCP2022_020269 | InChI=1S/C15H12N2O/c16-15(18)17-13-7-3-1-5-11(13)9-10-12-6-2-4-8-14(12)17/h1-10H,(H2,16,18) |

**Supplementary Table 3:** List of compounds selected from one screen from Curie Institut displayed in **Figure 3**

| ID in Fig. 5 | Metadata_JCP2022 | Metadata_InChI |
|---|---|---|
| C1 | JCP2022_090566 | InChI=1S/C13H14BrNO2/c1-3-4-5-15-11-8(2)6-9(14)7-10(11)12(16)13(15)17/h6-7H,3-5H2,1-2H3 |
| C2 | JCP2022_067845 | InChI=1S/C10H8BrNO2/c1-5-3-6(11)4-7-8(5)12(2)10(14)9(7)13/h3-4H,1-2H3 |
| C3 | JCP2022_078573 | InChI=1S/C13H14BrNO2/c1-7(2)6-15-11-8(3)4-9(14)5-10(11)12(16)13(15)17/h4-5,7H,6H2,1-3H3 |
| C4 | JCP2022_095748 | InChI=1S/C9H6N2O4/c1-4-2-5(11(14)15)3-6-7(4)10-9(13)8(6)12/h2-3H,1H3,(H,10,12,13) |
| C5 | JCP2022_016832 | InChI=1S/C9H7NO3/c1-13-5-2-3-7-6(4-5)8(11)9(12)10-7/h2-4H,1H3,(H,10,11,12) |
| C6 | JCP2022_016414 | InChI=1S/C9H6BrNO2/c1-4-2-3-5-6(7(4)10)8(12)9(13)11-5/h2-3H,1H3,(H,11,12,13) |
| C7 | JCP2022_043866 | InChI=1S/C9H7NO2/c1-5-3-2-4-6-7(5)8(11)9(12)10-6/h2-4H,1H3,(H,10,11,12) |
| C8 | JCP2022_103644 | InChI=1S/C8H4ClNO2/c9-4-1-2-6-5(3-4)7(11)8(12)10-6/h1-3H,(H,10,11,12) |
| C9 | JCP2022_042718 | InChI=1S/C8H5NO2/c10-7-5-3-1-2-4-6(5)9-8(7)11/h1-4H,(H,9,10,11) |
| C10 | JCP2022_091149 | InChI=1S/C9H7NO3/c1-13-6-4-2-3-5-7(6)10-9(12)8(5)11/h2-4H,1H3,(H,10,11,12) |
| C11 | JCP2022_057174 | InChI=1S/C9H4F3NO2/c10-9(11,12)5-3-1-2-4-6(5)13-8(15)7(4)14/h1-3H,(H,13,14,15) |

**Supplementary Table 4:** List of compounds from the chemical series displayed in **Figure 5**

| PubChem CID | Metadata_JCP2022 | Compound Name | Target | InChI |
|---|---|---|---|---|
| 9543416 | JCP2022_010404 | Adezamapimod | MAPK | InChI=1S/C21H16FN3OS/c1-27(26)18-8-4-16(5-9-18)21-24-19(14-2-6-17(22)7-3-14)20(25-21)15-10-12-23-13-11-15/h2-13H,1H3,(H,24,25)/t27-/m0/s1 |
| 8515 | JCP2022_046511 | SP600125 | MAPK8 | InChI=1S/C14H8N2O/c17-14-9-5-2-1-4-8(9)13-12-10(14)6-3-7-11(12)15-16-13/h1-7,17H |
| 5169 | JCP2022_073458 | SB-202190 | BRD4 | InChI=1S/C20H14FN3O/c21-16-5-1-13(2-6-16)18-19(14-9-11-22-12-10-14)24-20(23-18)15-3-7-17(25)8-4-15/h1-12,25H,(H,23,24) |
| 2051 | JCP2022_025157 | Tyrphostin AG 1478 | EGFR | InChI=1S/C16H14ClN3O2/c1-21-14-7-12-13(8-15(14)22-2)18-9-19-16(12)20-11-5-3-4-10(17)6-11/h3-9H,1-2H3,(H,18,19,20) |

**Supplementary Table 5:** List of compounds displayed in **Figure 6**
